# Whole genome sequencing provides novel insights into the evolutionary history and genetic adaptations of reindeer populations in northern Eurasia

**DOI:** 10.1101/2023.08.16.553162

**Authors:** Kisun Pokharel, Melak Weldenegodguad, Stephan Dudeck, Mervi Honkatukia, Heli Lindeberg, Nuccio Mazzullo, Antti Paasivaara, Jaana Peippo, Päivi Soppela, Florian Stammler, Juha Kantanen

## Abstract

Semi-domestic reindeer (*Rangifer tarandus tarandus*) play a vital role in the culture and livelihoods of indigenous people across the northern Eurasia. These animals are well adapted to harsh environmental conditions, such as extreme cold, limited feed availability and long migration distances. Therefore, understanding the genomics of reindeer is crucial for improving their management, conservation, and utilization. Here we have generated a new genome assembly for the Fennoscandian semi-domestic reindeer with high contiguity, making it the most complete reference genome for reindeer to date. The new genome assembly was utilized to explore genetic diversity, population structure and selective sweeps in Eurasian *Rangifer tarandus* populations which was based on the largest population genomic dataset for reindeer, encompassing 58 individuals from diverse populations. Phylogenetic analyses revealed distinct gene clusters, with the Finnish wild forest reindeer standing out as a unique sub-species. Divergence time estimates suggested a separation of ∼52,000 years ago between Northern-European *Rangifer tarandus fennicus* and *Rangifer tarandus tarandus*. Our study identified three main genetic clusters: Fennoscandian, the eastern/northern Russian and Alaskan group, and the Finnish forest reindeer. Furthermore, two independent reindeer domestication events were inferred suggesting separate origins for the semi-domestic Fennoscandian and eastern/northern Russian reindeer. Notably, shared genes under selection, including retroviral genes, point towards molecular domestication processes that aided adaptation of this species to diverse environments.

## Introduction

Reindeer (*Rangifer tarandus*) in the *Cervidae* family of ruminant mammals inhabits tundra and boreal forest regions in northern Eurasia and North America. Within *R. tarandus*, various ecotypes or often termed as subspecies have been identified, such as tundra reindeer (or mountain reindeer) (*R. t. tarandus*), North American caribou (*R. t. caribou*), forest reindeer (*R. t. fennicus*) and Arctic Svalbard reindeer (*R. t. platyrhynchus*), based on their biogeographic distributions, morphological characters and sedentary or migratory life-history strategies ^1^. In the Fennoscandia and the northernmost regions in Russia, the semi-domestic reindeer obviously descend from wild tundra reindeer while in more southern reindeer herding sites in Russia including southern longitudes in Siberian regions also forest reindeer are managed ^1^. The geographic distribution of wild and semi-domestic populations and tundra and forest reindeer populations overlaps in some regions, and there are reindeer herding cultures where the coexistence of wild and domestic populations has promoted intentional crossbreeding of domesticated and non-domesticated reindeer ^2^. Although hybridization can be an important source of genetic variation e.g., for adaptation, it may also have undesirable genetic effects on native phenotypes and fitness-related traits in wild populations ^3^. For example, hybrid animals between the wild forest reindeer and domestic tundra reindeer would not be accepted in the conservation of endangered wild forest reindeer in Finland.

Recently, the very first genetic *Rangifer*-studies applying genome sequence data and bioinformatic methods were published ^4–7^ and currently there are *de novo* genome assemblies available for Eurasian *R. t. tarandus* ^4,7^ and North American *R. t. caribou* ^6,8^. The recent *de novo* reference genomes of *R. t. tarandus* were assembled using Illumina technology sequence data ^4,7^. However, the traditional Illumina mate-pair libraries may not span all genomic repetitive elements resulting in less optimal, fragmented assemblies in terms of high number of scaffolds ^9,10^. To create less and longer scaffolds even at the sub-chromosomal level and to obtain a more contiguous assembly, chromosome conformation capture (3C) techniques, such as Hi-C and Chicago libraries are available ^11^. The improved *de novo* genome assembly is a critical tool in various genomics studies examining genome architecture, genomic diversity, associations between genomic and phenotypic data, evolutionary and population genomics ^8,10–12^. The 3C approaches can also be used to detect fundamental units of three-dimensional genomic architecture, such as topologically associated domains (TADs) which play an important role in the gene expression regulations ^13^.

The very first studies focusing on population genomics of *Rangifer* have been conducted applying the new *de novo* assemblies and whole-genome resequencing data to examine within- and between population diversity ^6–8^ the research issues of which have previously been investigated using mitochondrial DNA and autosomal microsatellites as genetic markers ^2,14,15^. Autosomal microsatellite, mitochondrial DNA data sets and whole genome resequencing have indicated relatively high genetic variation within semi-domestic reindeer populations which may be indicative of the early phase of reindeer domestication and breeding history and in few cases also can suggest events of introgression from wild populations ^2,7,15^. Genome data sets have provided novel and versatile insights into evolution, demographic history and spread of reindeer populations from refugia after the Last Glacial Maximum and effects of natural selection on several crucial genes, which have promoted the adaptation of reindeer to challenging northern environments ^5,7^. *PRDM9*-gene, for example, has been involved in recombination and speciation, *PRDM1* and *OPN4B* in retinal development and *GRIA1* in circadian rhythm ^7^.

In the present study, we promote the new research field of *Rangifer* genomics by describing our efforts to improve the current Fennoscandian reindeer reference genome ^7^ and publishing so far, the largest population genomics data on Eurasian *R. tarandus*. We used Chicago and Hi-C sequencing libraries and HiRise pipeline for the reindeer genome assembly to produce long scaffolds. We applied the improved assembly for investigations of genetic diversity, population structure and selective sweeps in North Eurasian domestic and wild reindeer populations. We have updated our resequencing data currently including genome sequences of 58 individuals and new sequence data of Fennoscandian (Finland), Nenets (Arkhangelsk in western Russia) and Even (Sakha Republic) semi-domestic reindeer and wild forest reindeer (*R. t. fennicus*) in Finland. We conducted the most comprehensive population genomic studies for reindeer populations from which several individuals were resequenced.

## Results

### 2.1. A new Fennoscandian reindeer reference genome

We generated a new reference genome assembly of the semi-domestic Fennoscandian reindeer derived from the same male reindeer individual the genome of which was sequenced to produce the first reference assembly including the mitochondrial genome ^7^. To obtain the present new reference assembly, we used these recent data based on the Illumina shotgun-sequencing technology and the present new sequence data obtained by sequencing Chicago and Hi-C proximity ligation libraries. The number and length of read pairs produced for the Chicago library 1 was 223 million and 2×150bp, respectively, and for the library 2 were 180 million and 2×150bp, respectively. Together, these Chicago library reads provided 93.85 X physical coverage of the genome (1-100 kb pairs). For the Hi-C proximity ligation libraries, the corresponding values were as follows: 203 million, 2×150bp for library 1; and 212 million, 2×150 bp for library 2. Together, these Dovetail Hi-C library reads provided 44,243.23 x physical coverage of the genome (10-10,000 kb pairs) The final assembly was comprised of 2,663.35 Mb with a total of 127,880 scaffolds of which 11,813 were greater than 1 kb. The longest scaffold in our assembly is 116,325,531 bp and the N50 and N90 scaffold lengths are 38.508 Mb (L50 = 15 scaffolds) and 69.739 Mb (L90 = 34 scaffolds), respectively (Table 1, Supplementary Figure 11). So far, five studies have reported genome assemblies for *R. tarandus* originating from different geographic regions (Table 1). In comparison with the metrics of existing assemblies, our new reference genome has fewer and longer scaffolds, thus indicating a good alternative reference for reindeer genomic studies. Moreover, synteny comparison showed that the top 37 scaffolds (all above 10 Mb) represent 95% of the reindeer assembly and covered the entire 30 (29 autosomes and Chr X) chromosomes of cattle reference genome (Figure 1). Cattle chromosomes 1, 2, 6, 8, 9, X split into two scaffolds each of reindeer whereas cattle chr 27, 28 were represented by one reindeer scaffold. There were a few non-syntenic regions, marked by intersecting lines/bands.

**Figure 1:**
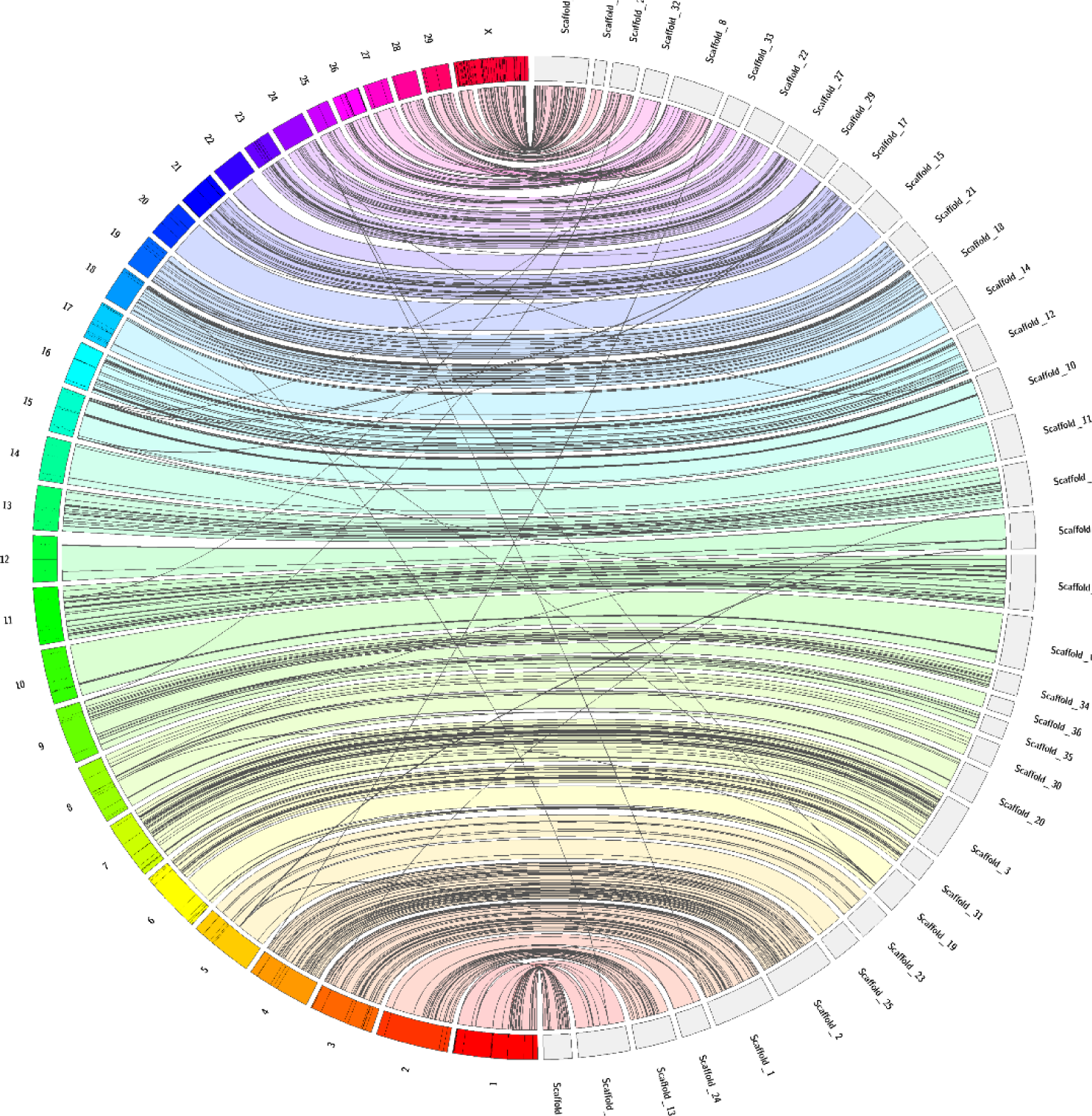
Jupiter consistency plot showing genome alignment between the reindeer assembly and cattle reference genome. Top 37 scaffolds (all above 10 Mb) represented 95% of the reindeer assembly and covered the entire 30 (29 autosomes and Chr X) chromosomes of cattle reference genome. Colored bands represent synteny between two genomes and the crossing lines indicated possible genomic rearrangements or break points in the scaffolds.

**Table 1:**
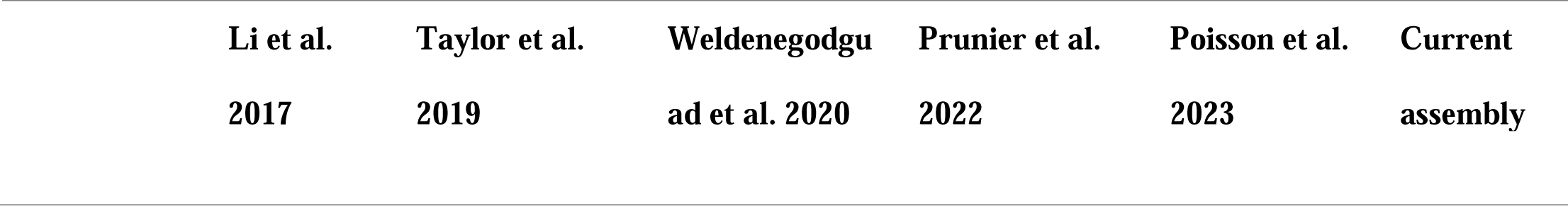
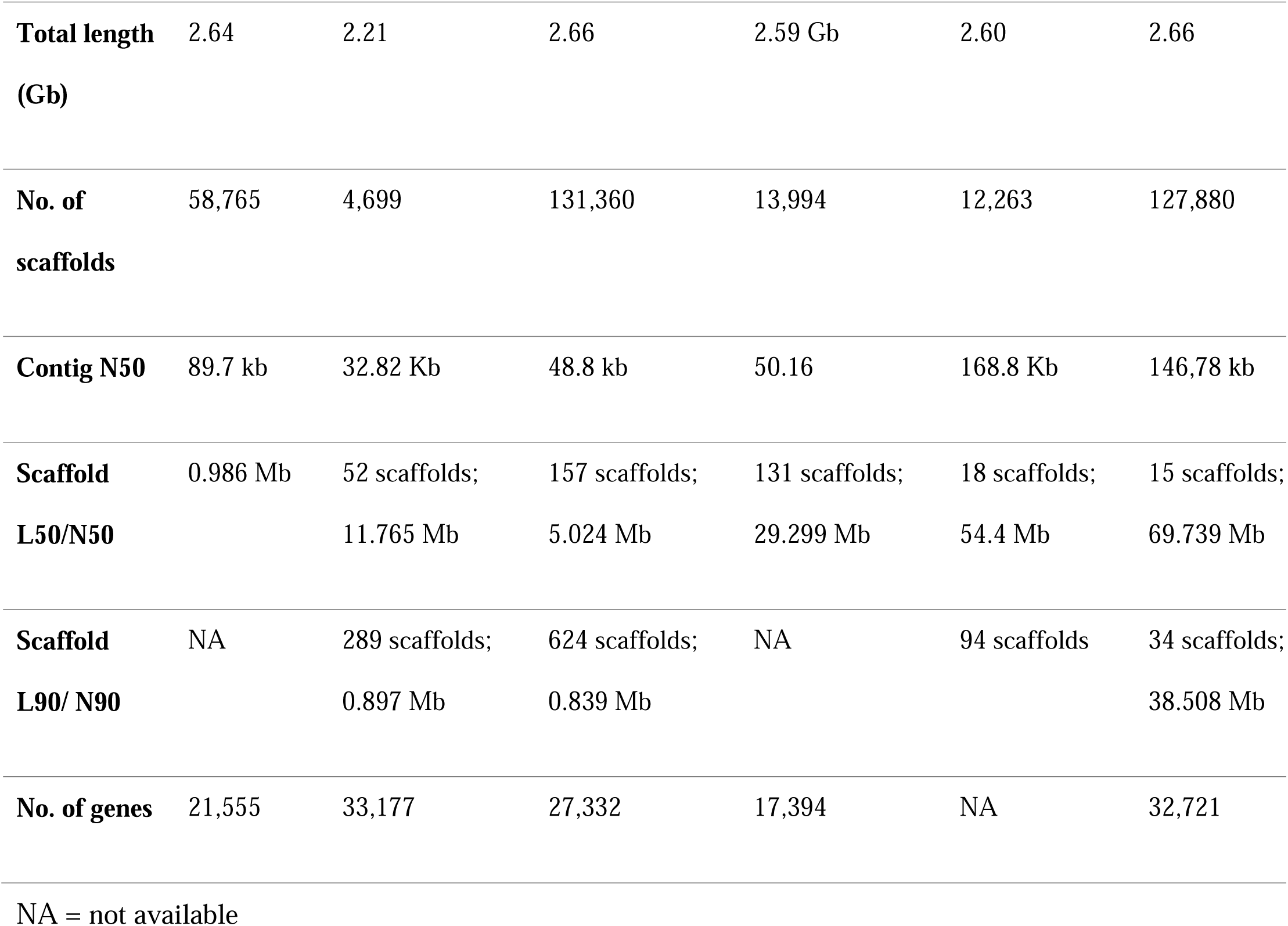
Comparative summary statistics of existing reindeer genome assemblies with our new assembly.

| Li et al. | Taylor et al. | Weldenegodgu | Prunier et al. | Poisson et al. | Current |
| --- | --- | --- | --- | --- | --- |
| 2017 | 2019 | ad et al. 2020 | 2022 | 2023 | assembly |

Reindeer population genomics
|  |  |  |  |  |  |  |
| --- | --- | --- | --- | --- | --- | --- |
| **Total length**<br>**(Gb)** | 2.64 | 2.21 | 2.66 | 2.59 Gb | 2.60 | 2.66 |
| **No. of**<br>**scaffolds** | 58,765 | 4,699 | 131,360 | 13,994 | 12,263 | 127,880 |
| **Contig N50** | 89.7 kb | 32.82 Kb | 48.8 kb | 50.16 | 168.8 Kb | 146,78 kb |
| **Scaffold**<br>**L50/N50** | 0.986 Mb | 52 scaffolds;<br>11.765 Mb | 157 scaffolds;<br>5.024 Mb | 131 scaffolds;<br>29.299 Mb | 18 scaffolds;<br>54.4 Mb | 15 scaffolds;<br>69.739 Mb |
| **Scaffold**<br>**L90/ N90** | NA | 289 scaffolds;<br>0.897 Mb | 624 scaffolds;<br>0.839 Mb | NA | 94 scaffolds | 34 scaffolds;<br>38.508 Mb |
| **No. of genes** | 21,555 | 33,177 | 27,332 | 17,394 | NA | 32,721 |

NA = not available

Genome contained 35.52% of repeats of which Class I transposable elements (TEs) repeats comprised of 30% and class II TEs repeats were 1.71% (TEs reviewed in eg. Lerat, 2010). Altogether 32,721 genes were present in our assembly with total coding region spanning 32,201,636 bp (% of genome) with average length of 984 base pairs. There were a total of 1694 single-exon genes. Benchmarking Universal Single Copy Orthologue (BUSCO) analysis ^17^ revealed 202 complete single-copy BUSCOs and 18.8% were missing. We manually curated 55 genes (Supplementary Data 1) of which seven could not be found.

Altogether 263 topologically associated domains (TADs) were detected at 10kbp resolution and with 50kbp resolution, we observed 1452 TADs (Supplementary Table 1). Those 263 TADs detected with the resolution of 10kbp resolution represented 6.44% of the genome with an average TAD size of 645,741bp. Moreover, the number of isochores and CCCTC-binding factor (CTCF) sites was 23,575 and 11,077, respectively.

### 2.2. Population genomic analysis

#### 2.2.1. Genomic variants

For the present study, 35 individuals of the Finnish, Nenets and Even semi-domestic reindeer and the wild Finnish Forest reindeer were resequenced (Supplementary Data 2). After the new raw resequenced data were processed, we generated a total of 1.57 Tb clean paired-end new data. In our previous *Rangifer* whole-genome data comprising of 23 animals ^7^ and pooled here to the present new data, we generated 680 Gb clean paired-end whole genome sequence (WGS) data. Alignment of the reads to the present assembled reference genome rta_v2.0 was successful, with 98.7% of the reads mapped to the reference genome on average per individual indicating that the qualities of whole genome sequence data were found to be su_Jiciently good for downstream analyses. The average sequencing coverage of the 58 genomes was 11.4 X, varying from 8.1 to 15.8 X coverage (Supplementary Data 2). Average depth was lower with slight variation in the older sample batch (NMBU-* samples), with newer sample batch having higher depth of coverage on average but greater variation in depth between the samples.

A total of 41.09 M high-quality SNPs were detected across all 58 individuals (the pooled data). The average number of SNPs detected per individual was 8.41 M (Supplementary Figure 3B). A total of 5.8 M indels was also detected across all samples, with average indel count per individual being 1.2 M (Supplementary Figure 3C). Of the 12 di_Jerent populations, the Even semi-domestic reindeer from the Sakha (Yakutia) Republic (9 animals) showed the highest number of SNPs, with 20.8 M SNPs found in total among the population and 9.38 SNPs detected on average per individual. At the other end of the spectrum, Svalbard wild arctic reindeer population (only 3 individuals) contained a total of 8.59 M SNPs, with average number of SNPs identified per individual being 7.17 M. Noteworthy is the split of the Nenets semi-domestic reindeer from Arkhangelsk into two groups, with SNP counts of 6.31-6.98 M in six individuals and SNP counts of 8.43-9.34 M in four individuals. Indel counts followed the same general trend, but on visual inspection of the plots, this trend appeared not to have a clear correlation with the sequencing depth. Also, the transition-to-transversion (TS/TV) ratios were uniform and within the expected range across all the samples (Supplementary Figure 4A; Supplementary Data 2) and populations (Supplementary Data 3), indicating consistent quality of the call set. The proportions of homozygous and heterozygous SNPs in Svalbard wild arctic reindeer population were clearly deviating from the other populations, showing a skew towards homozygous SNPs; this is in line with the Svalbard population being an isolated population with presumably high inbreeding (Supplementary Figure 4B).

#### 2.2.2. Genetic relationships between 59 animals

Genetic relationships between all the 59 animals including the reference animal were studied using principal component analysis (PCA). The PCA plot of the SNP data (Figure 2) displayed that the three Fennoscandian populations: Finnish semi-domestic reindeer, Norwegian semi-domestic reindeer and Norwegian wild tundra reindeer were in the same cluster, the Russian and Alaskan animals formed a more heterogenous cluster separated by the second principal component from the Fennoscandian populations and distinct clustering of the Svalbard wild arctic reindeer and Russian wild tundra reindeer from Novaja Zemlya based on the first principal component. Moreover, the Finnish wild forest reindeer were found in a distinct cluster separated by the second principal component from the two main clusters.

**Figure 2.**
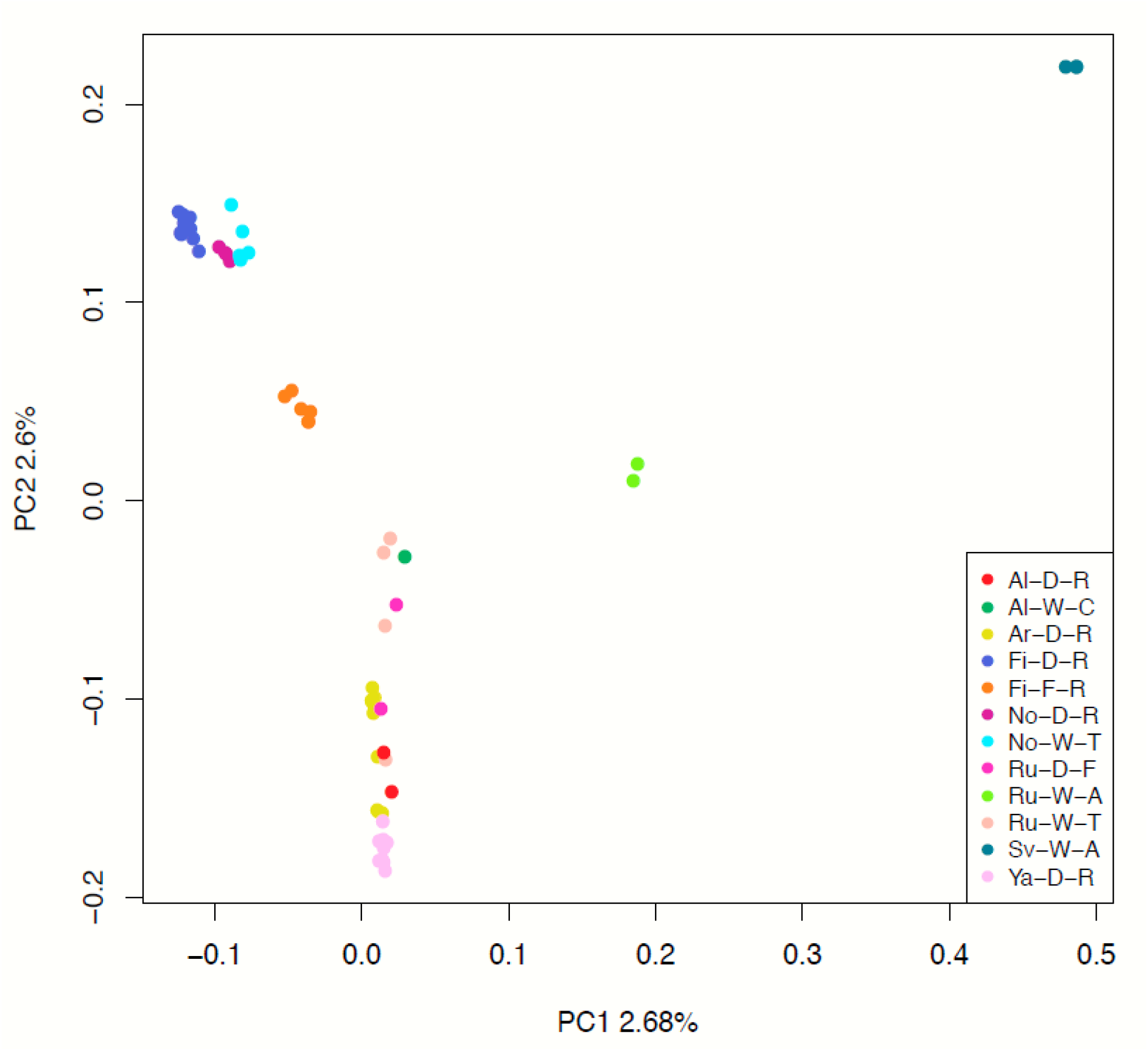
Principal component analysis (PCA) plot based on filtered and LD-pruned SNPs of all 59 samples. Population codes: Al-D-R – Alaskan semi-domestic reindeer; Al-W-C – Alaskan wild caribou; Ar-D-R – Nenets semi-domestic reindeer (Arkhangelsk); Fi-D-R - Finnish semi-domestic reindeer; Fi-F-R – Finnish wild forest reindeer; No-D-R - Norwegian semi-domestic tundra reindeer; No-W-T - Norwegian wild tundra reindeer; Ru-D-F - Russian semi-domestic forest reindeer; Ru-W-A - Russian wild tundra mountain reindeer (Novaja Zemlya); Ru-W-T - Russian wild tundra reindeer; Sv-W-A - Svalbard wild arctic reindeer; Ya-D-R - Even semi-domestic reindeer.

Genetic relationships of the individual reindeer were further studied by constructing a neighbor-joining (NJ) tree based on SNP data (Figure 3). As with PCA analysis, the populations formed distinct clusters. Two main phylogenetic clusters – the *Rangifer* of Fennoscandia and the *Rangifer* of the eastern/northern Russian Federation and Alaska - were identified with high Bootstrap confidence values. Within these major clusters, animals tended to group according to their know ancestries and geographic origins, such as the Nenets semi-domestic reindeer in Arkhangelsk, the Even semi-domestic reindeer in Sakha, the Finnish wild forest reindeer, the Norwegian semi-domestic reindeer, the Finnish semi-domestic reindeer and the Norwegian wild Tundra reindeer. Interestingly, in the phylogenetic tree the semi-domestic forest reindeer from the Zabaikal region in the southern Siberia (see Andersson et al. 2017^2^) grouped to the same branch with the Finnish wild forest reindeer with a high Bootstrap value. Moreover, the Svalbard wild arctic reindeer appeared clearly more distant to all the other populations.

**Figure 3:**
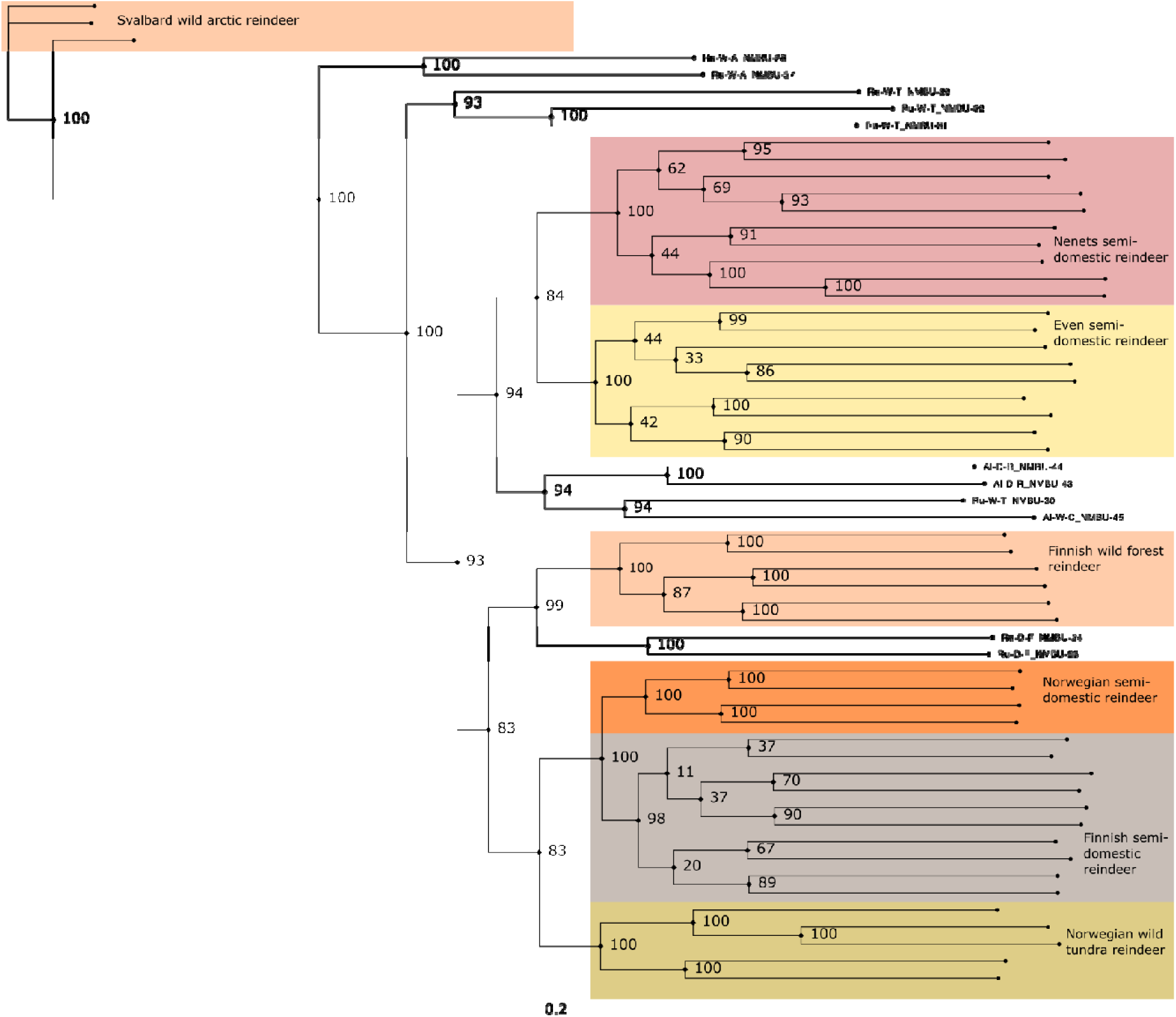
Genetic relationships between 59 animals. Neighbour-joining tree constructed showing genetic relationships between 59 animals calculated from the SNP data. Bootstrap confidence values obtained from 100 bootstrap replicates are shown at each branch. Seven main populations are highlighted in colors.

To infer population structure and admixture, ADMIXTURE software v.1.3 was used for population structure analysis including all 59 animals from 12 populations, using K values ranging from 2 to 12 (Figure 4). Cross-validation (CV) errors of the cluster numbers were lowest for K values 2, 3 and 4, increasing after that as K increased (Supplementary Figure 7). This and the PCA plot suggest that 4 would be the optimal number of ancestral populations. From the structure plots one can for example see that the Svalbard wild arctic reindeer are the only animals that are consistently assigned to their own unique cluster across all K (excluding K = 2 and 3) values. As indicated in the plot, K = 2 splits reindeer populations of Fennoscandian origin with that of Beringia refugial origin. At K = 3, the Finnish wild forest reindeer (Fi-F-R) showed distinctiveness from the Fennoscandian wild and semi-domestic tundra reindeer. At K = 4, the three Fennoscandian populations Fi-D-R, No-D-R and N - W-T were assigned to a separate cluster than the Even and Nenets semi-domestic reindeer while the Svalbard wild arctic reindeer and Finnish wild forest reindeer formed the two other clusters. Here, the two semi-domestic forest reindeer from the Zabaikal showed admixture of the Finnish wild forest reindeer and the Russian populations. In general, the clustering is well in line with the results of PCA and NJ tree results (Figures 2 and 3, respectively).

**Figure 4.**
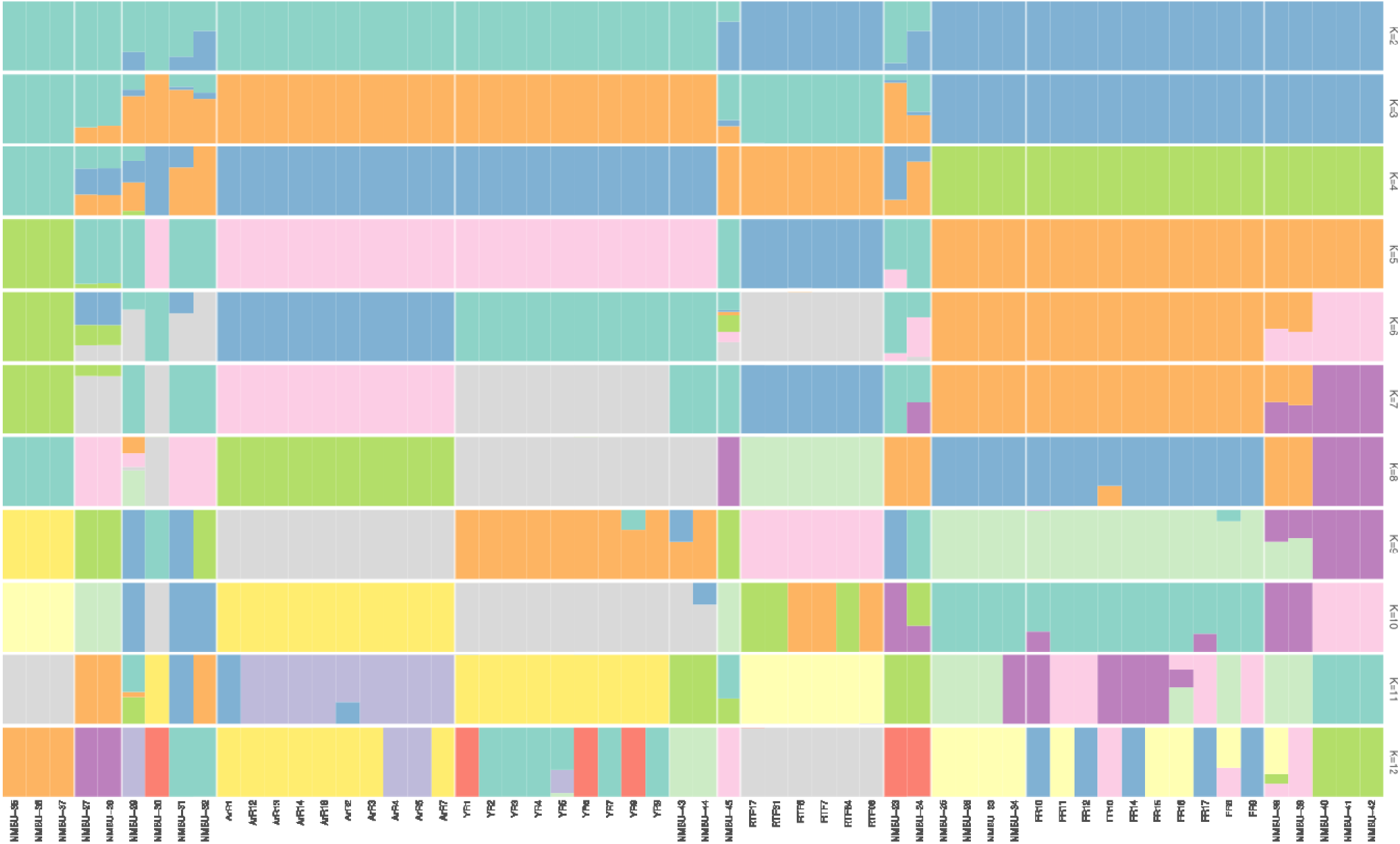
Population structure analysis of the twelve populations using ADMIXTURE. The bars represent individuals in a population and are segmented into colors based on the cluster assignment. The estimated proportion of the individual’s genome that belongs to a given cluster is indicated by the length of the colored segment. The analysis was repeated with different assumed numbers of clusters (K) that are indicated on the y-axis. Population codes and domestication status of the individuals are indicated on the x-axis. Individuals have been sorted within the population based on the cluster assignment values.

#### 2.2.3. Genetic analysis of seven *Rangifer*-populations

Based on the PCA, topology of the phylogenetic tree, geographic origins and the number of sequenced animals, seven “main” populations were selected for further within- and between-population genetic studies (see Table 2).

**Table 2.**
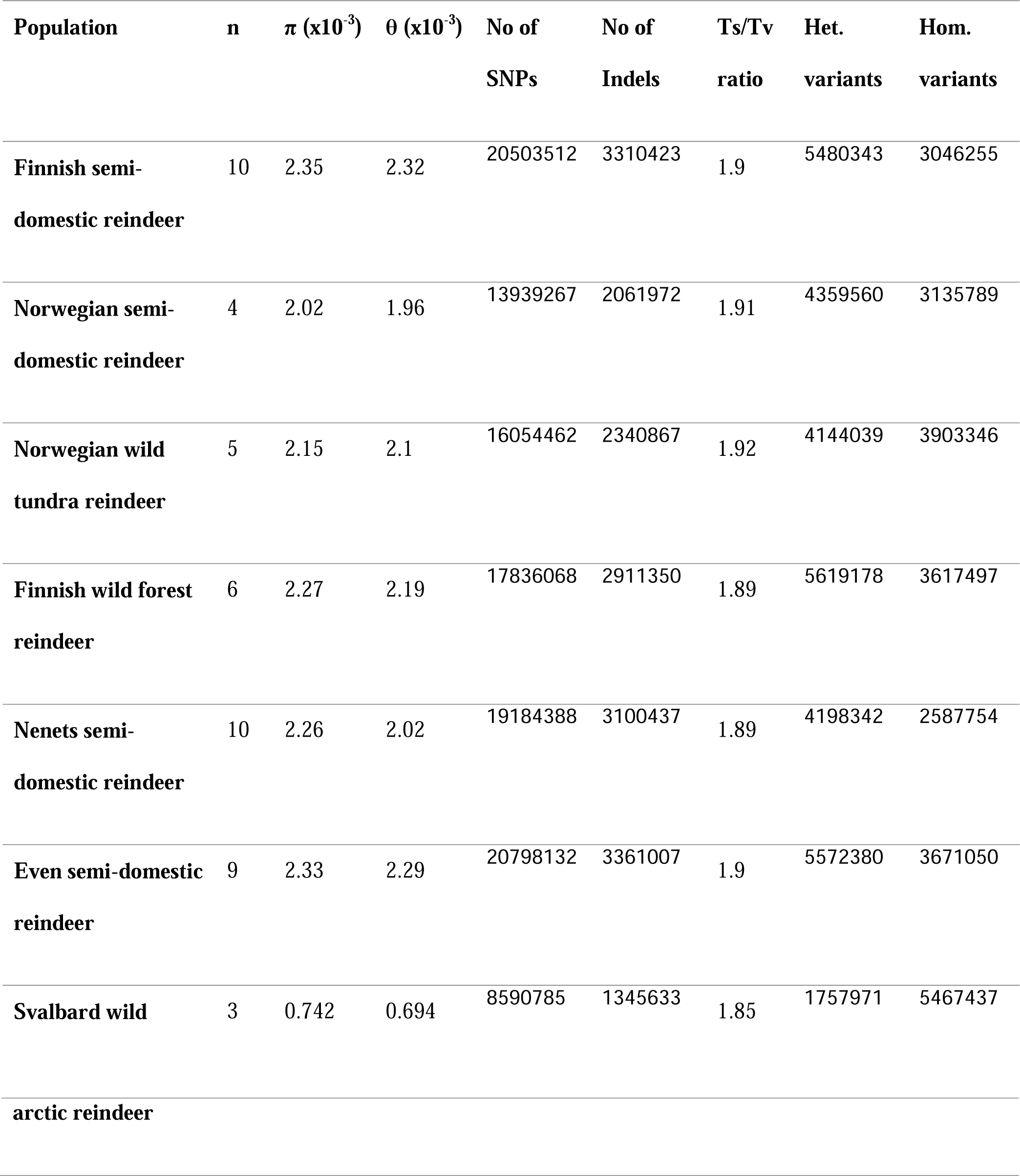
Population diversity statistics. Here, n represents the number of samples in each of the seven populations. Other statistics presented in the table are nucleotide diversity π, Watterson’s theta θ, number of SNPs and indels, transition to transversion (Ts/Tv) ratios as well as heterozygous and homozygous SNPs.

### 2.5. Population Diversity Statistics

Genetic diversity parameters were calculated for the reindeer populations based on the SNP data that was filtered with LD threshold 0.1 and MAF threshold 0.05 (totaling to 14.9M SNPs). The two main measures of overall genome-wide genetic diversity, pairwise nucleotide diversity π and expected level of diversity, Watterson’s theta θ, within the populations are shown in Table 2. The lowest π and θ values were, expectedly, in Svalbard population (0.742×10^-3^ and 0.694×10^-3^, respectively), while the highest values were found within the Finnish semi-domestic reindeer (2.35×10^-3^ and 2.32×10^-3^, respectively). Moreover, numbers of private (i.e. population-specific) variants were also compared between the populations. Svalbard reindeer and Norwegian semi-domestic reindeer had the smallest proportions of population-specific variants (only four animals were sequenced in the Norwegian reindeer), whereas the highest proportion of population-specific variants was found in Finnish semi-domestic reindeer (Table 2).

A population-level phylogenetic analysis was also conducted for the seven populations. For that, pairwise F_ST_ values were calculated and were used as the distance metric for building a NJ tree. The lowest pairwise F_ST_ was found between the Finnish and Norwegian semi-domestic reindeer, namely 0.009113, indicating a low diLerentiation between the populations (Supplementary Table 3). Highest F_ST_ values were found when Svalbard wild arctic reindeer were compared to other populations, the values ranging from 0.388949 (comparison to Even semi-domestic reindeer) to 0.424133 (comparison to Norwegian semi-domestic reindeer). As with the individual-level NJ tree (Supplementary Figure 6), the Svalbard population is clearly distinct from the other populations. There is also clear separation of the Even and Nenets semi-domestic reindeer populations (from Yakutia and Arkhangel, respectively) from the four Fennoscandian populations. F_ST_ values between one population and all others were also determined using a sliding window approach, dividing all analyzed SNP sites into 10 K SNP windows. Average F_ST_ values within each window were then plotted (Supplementary Figure 7).

### 2.7. Signatures of selection

In order to perform a genome-wide scans for selective sweeps, we focused on reindeer populations identified in the PCA (Figure 2) and phylogenetic (Figure 3) analysis and carried out selective sweep analysis using RAiSD software ^18^ in five sub-populations, namely Finnish wild forest reindeer, Norwegian wild tundra reindeer, Fennoscandian semi-domestic reindeer (the Finnish and Norwegian populations pooled), Nenets semi-domestic reindeer, and Even semi-domestic reindeer (Supplementary Data 4). We found many genomic regions exhibiting selective sweeps in each population - a total of 1538 in Finnish wild forest reindeer, 1352 in Norwegian wild tundra reindeer, 1928 in Fennoscandian semi-domestic reindeer, 1197 in Nenets semi-domestic reindeer and 1832 in Even semi-domestic reindeer – distributed in the top 40 scaffolds. These regions show significantly higher values of μ-statistics due to a result of positive selection, as μ-statistics is a measure of positive selection. The identified selective sweep genomic regions were mapped to several genes: 247 (Finnish wild forest reindeer), 290 (Norwegian wild tundra reindeer), 258 (Fennoscandian semi-domestic reindeer), 176 (Nenets semi-domestic reindeer) and 271 (Even semi-domestic reindeer) (Supplementary Data 4).

In our investigation, we found several genes related to cold adaptation, such as non-shivering thermogenesis, smooth muscle contraction, blood pressure, response to temperature, basal metabolic rate and energy metabolism^19^ which were under positive selection in Finnish wild forest reindeer (*CKMT2*, *EDN3* and *HSPB6*), Fennoscandian semi-domestic reindeer (*GCLM*), Nenets semi-domestic reindeer (*DNAJC1*, *DNAJC11* and *KCNB1*) and Even semi-domestic reindeer (*AHR*, *DNAJC11*, *GNAS* and *HR*). Moreover, the identified selective sweep genes in the populations include genes associated with immune response, for instance in Finnish wild forest reindeer (*LY9, FGB, TRAV12-3, TRAV14DV4, TRAV22* and *TRAV41*), Norwegian wild tundra reindeer (*ALOX15*, *CD3D*, *CD3G*, *ATF7*, *IFITM1*, *IFITM2*, *IFITM3*, *KLRC1*, *TGFB1*, *TRAV26-2*, *TRAV27*, *TRAV8-3*, *TRAV9-1* and *TRDV1*), Fennoscandian semi-domestic reindeer (*IRF8, MAP3K8, NFKB2, TRAV10, TRAV13-1, TRAV14DV4, TRAV26-2, TRAV8-3, TRDV1* and *TRIM5),* Nenets semi-domestic reindeer (*SEMA4D, TRAV22, TRAV24, TRAV26-1, TRAV26-2, TRAV27, TRAV8-3* and *TRDV1*) and Even semi-domestic reindeer (*AHR, ATRN, IFITM1, IFITM2, IFITM3, NLRP6, PLXNC1, TRAV12-3, TRAV14DV4, TRAV22, TRAV24, TRAV41* and *TRIM5*).

We also identified a number of genes under selection in the *Rangifer* population associated with ATP, lipid and energy metabolism, such as *APOO*, *CD5L, CD1E, ATP8B1, CKMT2, GABBR1* and *LIPK* in the Finnish wild forest reindeer, *CD5L, NUDT5, PERM1, P2RX7, SCP2* and *VPS9D1* in Norwegian wild tundra reindeer, *CD5L, CYP1B1, DNAH2, GNPAT, PERM1* and *SCP2* in Fennoscandian semi-domestic reindeer, *CD5L, FGGY, SCP2* and *SMCHD1* in Nenets semi-domestic reindeer and *APOM, ATP6V0A1, CD5L, PERM1* and *SCP2 in* Even semi-domestic reindeer.

In addition, we found several genes under positive selection associated with calcium binding, calcium metabolism and calcium homeostasis and circadian rhythm. Previous study has shown that genes associated with calcium metabolism and circadian rhythm are found to be associated with reindeer specific characteristics^5^. Examples of these genes associated with calcium binding, calcium metabolism and calcium homeostasis were *GABBR1*, *KCNIP1* and *NINL* in Finnish wild forest reindeer, *CACNA1E*, *CACNA2D2, GABBR1, PCDHA2, PCDHA9* and *SLC35G1* in Norwegian wild tundra reindeer, *OPRL1, PCDHA2, PCDHA5, PCDHA6, PCDHA7, PCDHA9, PCDHB15* and *VAPB* in Fennoscandian semi-domestic reindeer, *ADGRL2, PCDHA11, PCDHA5, PCDHA6, PCDHA7, PCDHB15* and *PCDHB18* in Nenets semi-domestic reindeer and *CDH15* in Even semi-domestic reindeer. Moreover, among the identified ‘selective sweep -genes’ there were genes know to be associated with circadian rhythm: *NFKB2* in Fennoscandian reindeer and *AHR*, *DRD4* and *HCRTR1 in* Even reindeer ^20–26^.

We looked for common genes under selection across all reindeer populations, as well as those unique to each of the five populations (Supplementary Data 5). As shown in the venn diagram (Figure 5) 18 genes were found to be under selection in all populations. These common genes included *Gag*, *Pol*, *RAB9A*, *MAN2A1*, *MTA3*, *sax-1* and *xpo1*. The highest number of genes under selection were present in Norwegian wild tundra reindeer on the other hand, Nenets semi-domestic reindeer had the lowest number of genes under selection.

**Figure 5:**
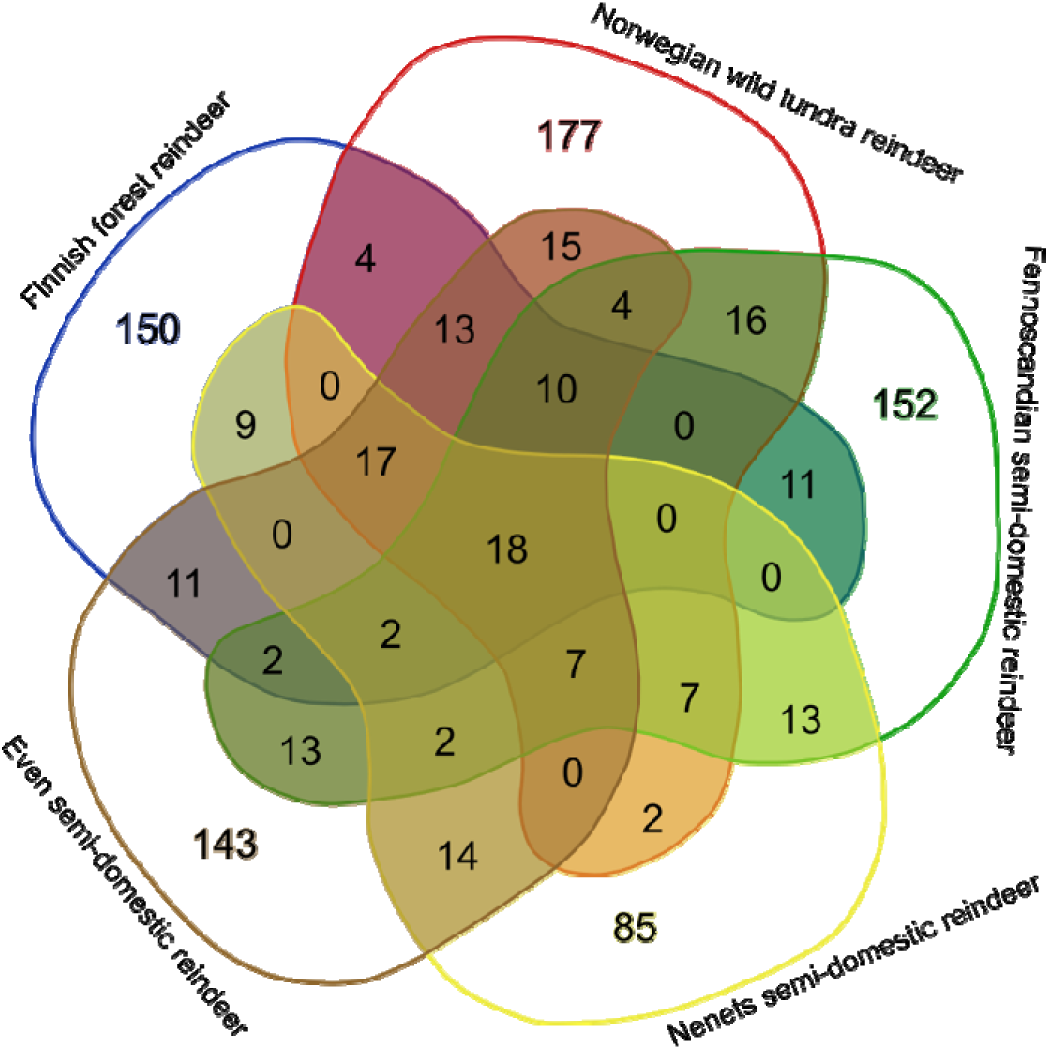
Distribution of genes under selection in five major groups: Finnish forest reindeer, Norwegian wild tundra reindeer, Fennoscandian semi-domestic reindeer, Nenets semi-domestic reindeer and Even semi-domestic reindeer.

### Divergence time estimations

We inferred the divergence time by coalescent hidden Markov model (CoalHMM) from Jocx tool for the following population pairs: Norwegian wild tundra *vs.* Finnish wild forest reindeer, Finnish wild forest reindeer *vs.* Even semi-domestic reindeer, Norwegian wild tundra *vs.* Even semi-domestic reindeer and Norwegian wild tundra reindeer *vs.* Norwegian semi-domestic reindeer using the top 39 scaffolds. Finnish wild forest reindeer was estimated to have diverged from Norwegian wild tundra and Even semi-domestic reindeer ∼51.6 thousand years ago (Kya) and ∼119.3Kya, respectively. Similarly, Norwegian wild tundra was estimated to have diverged from Norwegian semi-domestic reindeer and Even semi-domestic reindeer ∼12.0Kya and ∼22.0Kya, respectively. We found the divergence between Finnish wild forest reindeer and Even semi-domestic reindeer (∼119.3Kya) occurred earlier than between the Norwegian wild tundra and Even semi-domestic reindeer (22.0Kya).

## Discussion

We have generated and described here a highly contiguous new genome assembly for the Fennoscandian semi-domestic reindeer (*R. tarandus tarandus*) and used this updated assembly as a reference genome in the population genomic analyses. To date, five reference assemblies for reindeer/caribou have been published (Table 1). With N50 contig and N50 scaffold values of 146.78 kb and 69.739 Mb respectively, our reindeer assembly represents the most complete reference genome of reindeer. Pairwise genome comparison between our assembly with cattle reference genome indicated high synteny with 37 reindeer scaffolds mapping to all cattle chromosomes. All the 37 scaffolds are above 10 Mb in size and represent more than 95% of the assembly, and thus, our reference genome is assembled at near-chromosomal level. Here, we used the updated assembly to perform the most comprehensive population genomic study of the Eurasian reindeer species so far.

Our phylogenetic analyses (Figure 3) show that the Finnish forest reindeer (*R. t. fennicus*) is genetically distinct from the wild tundra and semi-domestic tundra reindeer in northern Europe. Based on the genetic distinctiveness and morphological and ecological differences found between the forest and tundra reindeer ^1^, we agree with the conclusion by Harding (2022) that the taxonomical status of ‘sub-species’ is pertinent for the Finnish Forest reindeer and tundra reindeer rather than that of two different ‘eco-types’. Moreover, we present here for the first time an estimate of the sub-species divergence time and found that the North-European *R. t. fennicus* and *R. t. tarandus* have diverged ∼52 000 years ago. This result indicates that the ‘sub-speciation’ began already before the ancestral populations of the present-day forest and tundra reindeer colonized North Europe after the Last Glacial Maximum, encompassing the period ca. 10 – 12 000 years ago ^27^. Furthermore, the origins of the ancestral populations spread to Northern Europe may have been in different refugia populations. Our studies on genetic structure of the northern Eurasian *Rangifer tarandus*-populations point towards three main genetic clusters suggesting the existence of three ancestral glacial refugia populations (Figure 2 and Figure 4): The Russian/northern American cluster reflecting the Euro-Beringian lineage, Fennoscandian cluster obviously descending from the South/Central European refugia populations ^7^ and the cluster of the Finnish forest reindeer. This conclusion of three main glacial refugia populations is in agreement with the partial mtDNA D-loop sequence analysis of 14 Eurasian and North American semi-domestic and wild *Rangifer tarandus* -populations ^14^. Interestingly, our phylogenetic and structure analyses showed genetic affinity between the wild Finnish forest reindeer and semi-domestic forest reindeer from Northeastern Zabaikal district, west from the Lake Baikal in southern Siberia. This novel finding suggests common ancestries of these two forest reindeer populations in a glacial refugium population. The wild forest reindeer became extinct in Finland in the beginning of the 1900’s, and the current population is based on animals which migrated back to eastern Finland from the Russian Karelia starting from 1950’s (Danilova et al. 2020). Archaeo-osteological evidences have indicated that the forest reindeer also originally spread to Finland from east ca. 7 500 years ago following the retracting ice margin while the tundra reindeer may have colonized the northern Fennoscandia especially along the Norwegian narrow ice-free coastal zone ^28^.

We found with our extended genomic data set (see also ^7^) that in both main geographic, genetically distinct population clusters (PCA, phylogenetic tree) there are semi-domestic animals suggesting at least two independent reindeer domestication events for Fennoscandian and Russian semi-domestic reindeer in the different parts of Eurasia. This conclusion is supported also by previous microsatellite and mtDNA studies ^29^. Our estimate suggests that the Fennoscandian and Russian (Yakutian Even reindeer analysed here) phylogenetic clusters diverged ∼22 000 years ago, while the Russian cluster and the wild Finnish forest reindeer cluster may have separated ∼119 000 years ago providing additional evidence for genetic distinctiveness of the Finnish forest reindeer in terms of its origin. Moreover, the divergence time estimations suggest that the Yakutian Even reindeer is taxonomically more like tundra reindeer sub-species type than that of the forest reindeer. Our estimate indicates that the divergence between the Norwegian wild tundra reindeer and the Norwegian semi-domestic reindeer occurred ∼12 000 years ago. However, ancestral wild reindeer populations of the Fennoscandian semi-domestic reindeer are not well known, and at least in the northernmost Fennoscandia local wild tundra reindeer may not have been domesticated when the Indigenous Sámi people shifted from reindeer-hunting based livelihood to reindeer pastoralism starting from the mid-16^th^ Century ^29,30^. These assumptions about the origins of the northern Fennoscandian semi-domestic reindeer are based on archaeological genetic studies on temporal changes and current distribution of common mtDNA haplogroups in semi-domestic and wild reindeer in Fennoscandia and Northwestern Russia (for review see ^29^). However, our whole-genome sequence data of individual animals of three Russian geographic populations and those of Nenets and Even reindeer do not reveal a candidate population whose ancestral population could have been the ancestral population for the Fennoscandian reindeer as well. However, interestingly, our genomic data indicated genetic affinity between the Nenets and Even reindeer from north-western (Arkhangelsk region) and north-eastern (Sakha Republic) Russia, respectively. This indicates that reindeer herding, and pastoralism have been based on animals of same genetic origin in a very large geographical area in the northern Russian territory among different northern indigenous societies. As pointed out by ^31^, these reindeer breeds belong to “Siberian reindeer” group of reindeer (*R. t. sibiricus*). In contrast to our finding, genotyping of limited number of autosomal microsatellites in the Russian reindeer breeds ^32,33^ clustered the Nenets and Even reindeer to different genetic sub-groups. The Nenets samples genotyped in the study by Svishchera et al. (2022) were collected in different regions (Yamal-Nenets and Khanty-Mansi regions) than the Nenets individuals sequenced here. Similarly, Kharzinova et al. (2020)^34^ found that the Nenets reindeer from the Arkhangelsk region was not closely related to the Even reindeer in Sakha ^34^. However, in that study, authors used Bovine HD BeadChip (Illumina) to characterize the various reindeer populations, the approach that is more prone to ascertainment bias.

In the phylogenetic treed based on genetic distances between the individuals (Figure 3), the Finnish wild forest reindeer individuals are grouped into two closely related branches supported by 100% bootstrap value. This shows that our samples of the Finnish Forest reindeer were from two geographical conservation areas, Kainuu and Suomenselkä. Moreover, our genomic data indicate that the individuals were not genetically influenced from the Finnish semi-domestic reindeer suggesting that the conservation of the Finnish wild forest reindeer has been successful as hybridization with semi-domestic reindeer may have not happened or it is rare.

Although the sample size, sequencing depth and the quality of a reference genome may effect on the level of genetic diversity seen in animal populations ^35^, the within-population genetic diversity estimates presented here (Table 2) indicate that the semi-domestic reindeer typically exhibit higher genetic diversity than, for example, domestic cattle breeds (*Bos taurus*) ^36^, domestic horse breeds (*Equus ferus caballus*) ^37^ and domestic sheep breeds (*Ovis aries*). Compared to domestic cattle, domestic horse and several other domesticated farm animal species, the semi-domestic reindeer is on the early stage of human-driven domestication. In addition to a less intensive human-made artificial selection, semi-domestic reindeer populations may have had a larger founder population sizes and possible admixture with wild reindeer populations could contribute to the level of within-population genetic diversity ^7^.

The genome-wide scans for selective sweeps conducted in this study focused on five distinct reindeer sub-populations: Finnish wild forest reindeer, Norwegian wild tundra reindeer, Fennoscandian semi-domestic reindeer (Finnish and Norwegian populations combined), Nenets semi-domestic reindeer, and Even semi-domestic reindeer. Mapping of the identified selective sweep genomic regions to specific genes showed distinct patterns in each population, with counts of genes under selection being 247, 290, 258, 176, and 271 for Finnish wild forest reindeer, Norwegian wild tundra reindeer, Fennoscandian semi-domestic reindeer, Nenets semi-domestic reindeer, and Even semi-domestic reindeer, respectively (Supplementary Data 4). The varying numbers of genes under selection in the different population groups highlight potential differences in adaptive pressures and genomic responses among these sub-populations. The genes identified under positive selection provide valuable insights into the adaptive processes shaping the genetic makeup of these reindeer populations. Notably, genes related to cold adaptation, non-shivering thermogenesis, smooth muscle contraction, blood pressure regulation, response to temperature, basal metabolic rate, and energy metabolism exhibited signs of positive selection in multiple reindeer populations ^19^. Additionally, immune response-related genes were highlighted, suggesting the importance of these genes in the context of local environmental challenges ^38^. Furthermore, genes associated with ATP, lipid, and energy metabolism were under selection, indicating the relevance of these metabolic pathways for reindeer survival and adaptation ^7,39^. Interestingly, genes linked to calcium binding, calcium metabolism, calcium homeostasis, and circadian rhythm also showed signals of positive selection. These findings align with previous studies linking these genes to reindeer-specific characteristics ^5^. These genes may play a pivotal role in maintaining physiological functions crucial for survival in the northern environments inhabited by reindeer. An investigation into common genes under selection across all reindeer populations identified 18 genes shared among all five sub-populations. These genes included *Gag*, *Pol*, *RAB9A*, *MAN2A1*, *MTA3*, *sax-1*, and *xpo1*. The presence of these shared genes suggests their significance in core adaptive processes across diverse reindeer populations. Interestingly, *gag* and *pol* are two of the three major proteins encoded within the retroviral genome^40^. These genes are considered to play important role in creating new gene families, the process which is defined as “molecular domestication”. Such phenomenon can help the organism adapt to new circumstances. The role of these genes and retrovirus in the adaptation and domestication of reindeer in the Northernmost Eurasia needs further research (see Chessa et al., 2009 ^41^). Our results indicate that the selection pressure for these genes were highest in the wild tundra and forest reindeer, and in Nenets and Even reindeer which are less managed by humans, compared to the Fennoscandian semi-domestic reindeer (also see Weldenegodguad et al., 2020, Supplementary Dataset 6).

## Conclusions

Here we have generated a highly contiguous genome assembly for the Fennoscandian semi-domestic reindeer (*Rangifer tarandus tarandus*) and utilized it as a reference genome for extensive population genomic analyses. The new assembly demonstrates high quality with contig and scaffold metrics indicating significant improvements compared to previous reindeer reference genomes. Phylogenetic analyses reveal genetic distinctiveness between Finnish wild forest reindeer and Norwegian wild tundra and semi-domestic reindeer in northern Europe. The genetic differentiation supports the idea that ‘sub-species’ is a more appropriate taxonomical classification for Finnish wild forest reindeer and Fennoscandian tundra reindeer than considering them as different ‘eco-types’. Notably, genetic affinity is detected between Finnish wild forest reindeer and semi-domestic forest reindeer from northeastern Siberia, implying shared ancestral populations. Moreover, genetic clusters of semi-domestic animals in both Fennoscandian and Russian populations suggest at least two independent reindeer domestication events for these regions. We identified here that genes related to retroviral elements (*gag* and *pol*) are identified as common genes under selection across the Fennoscandian reindeer populations all five groups. These genes play a role in "molecular domestication," potentially aiding adaptation to new circumstances. These genes show stronger selection pressure in less managed populations and wild tundra/forest reindeer. Overall, the study provides insights into the complex evolutionary history, domestication, and genetic adaptation of reindeer populations across different regions. It sheds light on the genetic basis of adaptations related to climate, environment, and human interaction, opening avenues for further research into the unique features of reindeer in the northernmost parts of Eurasia.

## Materials and Methods

### Sample information

Animal handling procedures and sample collections were performed in accordance with the legislations approved by the Animal Experiment Board in Finland (ESAVI/7034/04.10.05.2015) and the Russian Authorization Board (FS/U.VN-03/163733/07.04.2016). In our previous study^7^, we used the Illumina technology and assembled *de novo* the genome of one-year-old male reindeer (*R. tarandus tarandus*) from Sodankylä, Finland. In the present study, we improved the genome assembly of this same individual. DNA for library preparations and sequencing was extracted using a standard phenol-chloroform extraction from liver and muscle samples which were collected at slaughter and stored in stored in RNA*later*® Solution (Ambion/QIAGEN, Valencia, CA, USA).

For the resequencing and population genome analyses, we collected samples from Fennoscandian reindeer of Muddusjärvi and Sallivaara herding cooperatives in northern Finland (n=10 males, blood samples in EDTA tubes), Nenets reindeer from the Arkhangelsk region in North-West Russia (n=2 females and 8 males, hair samples), Even reindeer from Eveno-Bytantay region, northern Sakha, Russia (n=3 females and 6 males, blood samples in EDTA tubes) and wild forest reindeer (*R. t. fennicus*) in Finland (n=5 females and 1 male, blood samples in EDTA tubes). DNA was extracted using DNeasy Blood and Tissue Kits (Qiagen, Valencia, CA, USA). In addition, we included recently published data on 23 *Rangifer* genomes^7^ in this study: semi-domestic forest reindeer from Russia (n=2), semi-domestic tundra reindeer from Norway (n=4), wild tundra reindeer from Russia (n=2), wild tundra reindeer from Russia (n=4), wild tundra reindeer from Norway (n=5), wild arctic reindeer from Svalbard, Norway (n=3), Alaskan semi-domestic reindeer from USA (n=2) and Alaskan wild caribou from USA (n=1). More detailed information of these animals and DNA extraction of the samples are given in Flagstad and Røed (2003)^14^.

### Library preparation and sequencing

Four Chicago and HiC libraries were prepared ^9^ in the Dovetail laboratory (www.dovetailgenomics.com). For each library, ∼500ng of gDNA (mean fragment length = 70 kb) was reconstituted into chromatin *in vitro* and fixed with formaldehyde. Fixed chromatin was digested with DpnII, the 5’ overhangs filled in with biotinylated nucleotides, and then free blunt ends were ligated. After ligation, crosslinks were reversed, and the DNA purified from protein. Purified DNA was treated to remove biotin that was not internal to ligated fragments. The DNA was then sheared to ∼350 bp mean fragment size and sequencing libraries were generated using NEBNext Ultra enzymes and Illumina-compatible adapters. Biotin-containing fragments were isolated using streptavidin beads before PCR enrichment of each library. The libraries were sequenced on an Illumina HiSeq X platform.

Two Dovetail HiC libraries were prepared in a comparable manner as described previously ^42^. For each library, chromatin was fixed in place with formaldehyde in the nucleus and then extracted. Fixed chromatin was digested with DpnII, the 5’ overhangs filled in with biotinylated nucleotides, and then free blunt ends were ligated. After ligation, crosslinks were reversed, and the DNA purified from protein. Purified DNA was treated to remove biotin that was not internal to ligated fragments. The DNA was then sheared to ∼350 bp mean fragment size and sequencing libraries were generated using NEBNext Ultra enzymes and Illumina-compatible adapters. Biotin-containing fragments were isolated using streptavidin beads before PCR enrichment of each library. The libraries were sequenced on an Illumina HiSeq X platform.

The input *de novo* assembly, shotgun reads, Chicago library reads, and Dovetail HiC library reads were used as input data for HiRise, a software pipeline designed specifically for using proximity ligation data to scaffold genome assemblies ^9^. An iterative analysis was conducted. First, Shotgun and Chicago library sequences were aligned to the draft input assembly using a modified SNAP read mapper ^43^. The separations of Chicago read pairs mapped within draft scaffolds were analyzed by HiRise to produce a likelihood model for genomic distance between read pairs, and the model was used to identify and break putative mis-joins, to score prospective joins, and make joins above a threshold. After aligning and scaffolding Chicago data, Dovetail HiC library sequences were aligned and scaffolded following the same method. After scaffolding, shotgun sequences were used to close gaps between contigs.

### TAD Analysis

Hi-C contact matrices in two formats, namely cool and hic, were generated. Both contact matrices were generated from the same BAM file by using read pairs where both ends were aligned with a mapping quality of 60. TADs were identified using the Arrowhead program implemented in the Juicertools package ^44^. We call TADs at 3 different resolutions: 10 kbp, 25 kbp, and 50 kbp. The parameters used were -k KR -m 2000 -r 10000, -k KR -m 2000 -r 25000, and -k KR -m 2000 -r 50000. A/B compartments were identified at 1 Mbp using the eigenvector program implemented in the Juicertools package. The parameters used were KR BP 1000000. Isochores were predicted using the isofinder program ^45^. The parameters used were 0.90 p2 3000. The output was post-processed to convert it to a bedpe format. CTCF sites were predicted using the CREAD program ^46^. The position weight matrix was downloaded from CTCFBSDB 2.0 website. The output was then post-processed to convert it to a bed file. Multires files were generated using clodius package. These files can be loaded in HiGlass: an open-source visualization tool ^47^.

### Genome Annotation

Repeat families found in the genome assemblies of *Rangifer tarandus* were identified *de novo* and classified using the software package RepeatModeler (version 2.0.1) (http://www.repeatmasker.org/RepeatModeler/). RepeatModeler depends on the programs RECON (version 1.08) and RepeatScout (version 1.0.6) for the de novo identification of repeats within the genome. The custom repeat library obtained from RepeatModeler were used to discover, identify and mask the repeats in the assembly file using RepeatMasker (Version 4.1.0)^48^.

Coding sequences from *Bos taurus*, *Rangifer* tarandus (http://www.caribougenome.ca/downloads) ^6^, *Rangifer tarandus* (chinese - http://animal.nwsuaf.edu.cn)^4^, *Ovis aries* and *Capra hircus* were used to train the initial ab initio model for *Rangifer tarandus* using the AUGUSTUS software (version 2.5.5) ^49^. Six rounds of prediction optimization was done with the software package provided by AUGUSTUS. The same coding sequences were also used to train a separate ab initio model for *Rangifer tarandus* using SNAP (version 2006-07-28) ^50^. RNA-Seq data from four adipose tissues were used for improiving the annotation. RNAseq reads were mapped onto the genome using the Spliced Transcripts Alignment to a Reference (STAR) aligner software (version 2.7) ^51^ and intron hints generated with the bam2hints tools within the AUGUSTUS software. MAKER ^52,53^ SNAP, and AUGUSTUS (with intron-exon boundary hints provided from RNA-Seq) were then used to predict genes in the repeat-masked reference genome. To help guide the prediction process, Swiss-Prot peptide sequences from the UniProt database were downloaded and used in conjunction with the protein sequences from *Bos taurus*, *Rangifer tarandus* (http://www.caribougenome.ca/downloads), *Rangifer tarandus* (Chinese - http://animal.nwsuaf.edu.cn/), *Ovis aries* and *Capra hircus* to generate peptide evidence in the Maker pipeline. Only genes that were predicted by both SNAP and AUGUSTUS softwares were retained in the final gene sets. To help assess the quality of the gene prediction, AED scores were generated for each of the predicted genes as part of the MAKER pipeline. Genes were further characterized for their putative function by performing a BLAST search of the peptide sequences against the UniProt database. tRNA sequences were predicted using the software tRNAscan-SE (version 2.05) ^54^.

### Population genomic data analysis

For the various population genomic analyses, we have whole-genome sequencing data of 58 *Rangifer*-individuals (Supplementary Data 2): 23 genomes from our previous study ^7^ and a new set of 35 individuals sequenced in the present study. Whole-genome sequencing of DNA samples of these new individuals was done in BGI as previously described ^7^. The whole data set (58 genomes) was used only in the Principal Component Analysis (PCA) and in the individual based phylogenetic analysis, while for more comprehensive population genomic analyses were done for 7 populations in which genomes of several individuals were sequenced. These populations were Finnish semi-domestic reindeer (Fi-D-R, n=10), Norwegian semi-domestic reindeer (No-D-R, n=4), Norwegian wild tundra reindeer (No-W-T, n=5), Finnish wild forest reindeer (Fi-F-R, n=6), Nenets semi-domestic reindeer from the Arkhangelsk region (Ar-D-R, n=10), Even semi-domestic reindeer from Sakha (Yakutia) (Ya-D-R, n=9) and wild arctic Svalbard reindeer (Sv-W-A, n=3). Our new and improved reindeer genome assembly was used as a reference in the population genomic analyses.

### QC and preprocessing

The quality of the raw read data was inspected using FastQC software, v. 0.11.8 ^55^. MultiQC, v1.8.dev0 ^56^ was used for generating quality control reports of all samples.

### Alignment

Samples were aligned with BWA, v. 0.7.17-r1188 ^57^ against the assembled reindeer reference genome reindeer using default parameters. Before the alignment, the contig names in the reference were modified to be in line with SAM format. Resulting bam files were sorted and indexed using SAMTools, v. 1.9 ^58^. Duplicate alignments were marked with PicardTools, v. 2.18.16 (https://broadinstitute.github.io/picard/)

### Variant Calling

SNPs and indels were called according to the GATK best practice guidelines using GATK v. 4.0.11^59^. First, HaplotypeCaller was used for calling variant from individual duplicate-marked alignment files. Per-sample gVCF files produced by HaplotypeCaller were combined into a multisample gVCF file using CombineGVCFs tool. GenotypeGVCFs tool was then used for joint genotyping of all samples. Separate SNP and indel vcf files were generated for plotting of quality scores to select appropriated thresholds for hard filtering of the variants. Based on the plots and the recommendations in GATK 4 User Guide, the following filters were applied to variant set: FS > 60.0, MQ < 40.0, MQRankSum < -8.0, QD < 2.0, ReadPosRankSum < -8.0 and SOR > 3.0 for SNPs and FS > 200.0, MQ < 40.0, QD < 2.0, ReadPosRankSum < -8.0 and SOR > 5.0 for indels. Variants that passed all filters were extracted using SelectVariants tool to create the final high-quality set of variants. Variant call statistics were generated using bcftools, v. 1.9 ^60^ and MultiQC, v1.8.dev0 ^56^.

### Principal Component analysis

Principal component analysis (PCA) was performed using R ^61^ package SNPRelate, v. 1.18.1 ^62^ using the filtered variant data as input. Linkage disequilibrium (LD)-based pruning using snpgdsLDpruning function was first applied to avoid strong influence of linked SNP clusters. Two diLerent LD thresholds, 0.2 and 0.5 were used for filtering the SNP set. snpgdsPCA function was then utilised for plotting the PCA results.

In an alternative approach, genotype probabilities rather than called genotypes were calculated using ANGSD software, v. 0.929 ^63^ using the following parameters: -uniqueOnly 1 - remove_bads 1 - only_proper_pairs 1 -trim 0 -C 50 -baq 1 -minMapQ 20 -minQ 20 -minInd 29 -setMinDepth 5 -setMaxDepth 100 -doCounts 1 -GL 1 -doMajorMinor 1 -doMaf 1 -skipTriallelic 1 -SNP_pval 1e-3 - doGeno 8 -doPost 1. 254891 sites were retrieved and used for calculating pairwise genetic distance using ngsDist from ngsTools, v. 1.0.2 ^64^.

### Population Diversity statistics

Population diversity statistics were calculated from the SNP data using R package PopGenome, v. ^65^. The analyzed SNP data consisted of the SNPhylo-filtered set of SNPs as described in ‘Phylogenetic Analysis’ section. For the seven main populations (Fi-D-R, No-D-R, No-W-T, Fi-F-R, Ar-D-R, Ya-D-R and Sv-W-A), the average pairwise nucleotide diversity within a population (π) and the proportion of polymorphic sites (Watterson’s *θ*) were calculated from the SNP data using the Bio::PopGen::Statistics package in BioPerl (v1.6.924) ^66^. Moreover, average minor allele frequencies of SNPs were calculated for those seven main populations in addition to counts of population-specific, private SNPs. FST and πi values were also calculated for consecutive windows of 50 K SNPs and average values of SNPs within the windows were plotted. Similar analysis of site frequency spectrum (SFS) values was conducted.

### Phylogenetic analysis

Neighbor-Joining tree of all samples was generated from the SNP data using SNPhylo, v. 20160204 ^67^. First, the SNPs were further filtered using SNPhylo using the following filter thresholds: Minimum depth of coverage > 5, the percent of low-coverage samples < 5%, the percent of samples with no SNP information < 5%, LD < 0.1 and minor allele frequency (MAF) > 0.05. SNPhylo-generated sequences from the SNP data and performed multiple alignment of the sequences. PHYLIP tools^68^ were then used for computing a protein distance matrix and creating a neighbor-joining tree. FigTree v. 1.4.4 ^69^ was used for enhancing the tree appearance. A neighbor-joining tree with bootstrap values was generated using PHYML 3.0 software, using the multiple sequence alignments from SNPhylo as input. 100 bootstrap replicates were generated, and the obtained bootstrap tree was exported in newick format to FigTree that was again used for enhancing the tree appearance and adding the bootstrap support values to branches.

For population-level neighbour-joining tree, average pairwise F_ST_ values of the seven main populations (Fi-D-R, No-D-R, No-W-T, Fi-F-R, Ar-D-R, Ya-D-R and Sv-W-A) were computed using vcftools, v. 0.1.13 ^70^. Distance matrix was generated of the pairwise values, and the matrix was used for building a neighbour-joining tree using MEGA7 software ^71^.

### Population structure analysis

Population admixture analysis was performed using ADMIXTURE software, v. 1.3 ^72^. Hard-filtered SNP data was first converted into binary PLINK format using PLINK, v. 1.07 ^73^, after which 500 000 randomly sampled SNPs were extracted for the analysis. ADMIXTURE was then run using different K values ranging from 2 to 12 and with bootstrap parameter set to 200 replicates for estimation of standard errors. Population structure plots were generated using R package pop helper^74^.

### Positive selective sweep analysis

Genomic scans for positive selective sweeps were performed using RAiSD v 2.9 ^18^ using default parameters. RAiSD calculates the μ-statistic, a composite evaluation test that scores genomic locations by quantifying changes in the SFS, the levels of LD, and the amount the amount of genetic variation along the chromosome ^18^. In the analysis, we pooled the data sets of the two Fennoscandian semi-domestic reindeer populations (the Finnish and Norwegian reindeer) to improve the statistical power of the selective sweep analysis. Our PCA and phylogenetic analyses (see Figures 2 and 3) showed close genetic affinity between these reindeer populations inhabiting similar biogeographic northern regions. We performed a selective sweep analysis for each of five subpopulations (wild Finnish forest reindeer, wild Norwegian tundra reindeer, Fennoscandian semi-domestic reindeer, Nenets semi-domestic reindeer, and Even semi-domestic reindeer) (Supplementary Data 4). We selected scaffold length > 9 Mb (top 40 scaffold) for selective sweep analysis and RAiSD was run separately for each scaffold. To detect highly supported sweeps, we focused on each scaffold with top 1% of the μ-statistic. The cutoff value for μ-statistics was taken as the 99.99 percentile of the empirical distribution across the genome for each scaffold. Finally, the outliers selective sweep regions were manually annotated with the reference annotation (.gtf) file using bedtools v2.29.0 ^75^.

### Estimation of split time between different reindeer ecotypes under the CoalHMM Framework

The divergence times between different reindeer ecotypes were estimated using the coalescent hidden Markov models (CoalHMM) implemented in Jocx ^76^. To estimate the split time, we selected tundra and forest reindeer ecotypes from four main clusters based on the PCA plot (Figure 2): (Fi-F-R, No-W-T, No-D-R and Ya-D-R) and estimated divergence times between the population pairs: No-W-T *vs.* Fi-F-R, Fi-F-R *vs.* Ya-D-R, No-W-T *vs.* Ya-D-R and No-W-T *vs.* No-D-R. Fi-F-R. For the analysis, we selected a sample with the highest coverage from each population (NMBU-33, NMBU-39, RTF96 and YR6) and using the BAM mapping files generated a consensus pseudo-genome for each sample using ANGSD v0.931 ^63^ (*-doFasta 3*). In ANGSD we used *(-minMapQ 30*) to discarded reads with a minimum mapping quality lower than 30. Additional filter parameters (*-uniqueOnly 1*) was also used to discarded reads that doesn’t map uniquely. We split the pseudo-genomes into non-overlapping 10Mbp segments using seqkit ^77^. In each genome, we identified a total of 232 non-overlapping 10Mbp segments across the top 39 scaffolds (scaffold length greater than 10Mb). Maximum-likelihood estimation of the divergence time was then performed independently based on each 10Mbp segment using isolation-model from Jocx tool (https://github.com/jade-cheng/Jocx). The demographic model we used in this analysis is an isolation-model without gene flow after split. We used a mammalian mutation rate of 2.2 × 10^−9^ per base per year ^78^ to rescale the estimated model parameter to years.

## Acknowledgements

This study was funded by the Academy of Finland in the Arctic Research Programme ARKTIKO (decision number 286040). The authors thank Tiina Reilas, Innokentyi Ammosov and Knut H. Røed for the collaboration in collection of research materials, and reindeer herders for providing access to their reindeer for blood sample collection. We thank CSC-IT Center for Science, Finland for computational resources.

## Author contributions

J.K. designed the study. S.D., M.H., H.L., N.M., A.P., J.P., P.S., and F.S. collected the samples. K.P. and M.W. were involved in the data analysis. J.K., K.P., and M.W., interpreted the results. K.P. wrote the manuscript with contributions from J.K. and M.W. All authors read and approved the final manuscript.

## References

1. Pelletier, M., Kotiaho, A., Niinimäki, S. & Salmi, A.-K. Identifying early stages of reindeer domestication in the archaeological record: a 3D morphological investigation on forelimb bones of modern populations from Fennoscandia. Archaeol. Anthropol. Sci. 12, 169 (2020).

2. Anderson, D. G., Kvie, K. S., Davydov, V. N. & Røed, K. H. Maintaining genetic integrity of coexisting wild and domestic populations: Genetic differentiation between wild and domestic Rangifer with long traditions of intentional interbreeding. Ecol. Evol. 7, 6790–6802 (2017).

3. Iacolina, L., Corlatti, L., Buzan, E., Safner, T. & Šprem, N. Hybridisation in European ungulates: an overview of the current status, causes, and consequences. Mammal Rev. 49, 45–59 (2019).

4. Li, Z. et al. Draft genome of the reindeer (Rangifer tarandus). GigaScience 6, (2017).

5. Lin, Z. et al. Biological adaptations in the Arctic cervid, the reindeer (Rangifer tarandus). Science 364, eaav6312 (2019).

6. Taylor, R. S. et al. The Caribou (Rangifer tarandus) Genome. Genes 10, 540 (2019).

7. Weldenegodguad, M. et al. Genome sequence and comparative analysis of reindeer (Rangifer tarandus) in northern Eurasia. Sci. Rep. 10, 8980 (2020).

8. Prunier, J. et al. CNVs with adaptive potential in Rangifer tarandus: genome architecture and new annotated assembly. Life Sci. Alliance 5, e202101207 (2022).

9. Putnam, N. H. et al. Chromosome-scale shotgun assembly using an in vitro method for long-range linkage. Genome Res. (2016) doi:10.1101/gr.193474.115.

10. Elbers, J. P. et al. Improving Illumina assemblies with Hi-C and long reads: An example with the North African dromedary. Mol. Ecol. Resour. 19, 1015–1026 (2019).

11. Renaud, G. et al. Improved de novo genomic assembly for the domestic donkey. Sci. Adv. 4, eaaq0392 (2018).

12. Giani, A. M., Gallo, G. R., Gianfranceschi, L. & Formenti, G. Long walk to genomics: History and current approaches to genome sequencing and assembly. Comput. Struct. Biotechnol. J. 18, 9–19 (2020).

13. Han, J., Zhang, Z. & Wang, K. 3C and 3C-based techniques: the powerful tools for spatial genome organization deciphering. Mol. Cytogenet. 11, 21 (2018).

14. Flagstad, Ø. & Røed, K. H. Refugial origins of Reindeer (Rangifer tarandusL.) inferred from mitochondrial DNA sequences. Evolution 57, 658–670 (2003).

15. Røed, K. H. et al. Genetic analyses reveal independent domestication origins of Eurasian reindeer. Proc. R. Soc. B Biol. Sci. 275, 1849–1855 (2008).

16. Lerat, E. Identifying repeats and transposable elements in sequenced genomes: how to find your way through the dense forest of programs. Heredity 104, 520–533 (2010).

17. Manni, M., Berkeley, M. R., Seppey, M. & Zdobnov, E. M. BUSCO: Assessing Genomic Data Quality and Beyond. Curr. Protoc. 1, e323 (2021).

18. Alachiotis, N. & Pavlidis, P. RAiSD detects positive selection based on multiple signatures of a selective sweep and SNP vectors. Commun. Biol. 1, 1–11 (2018).

19. Cardona, A. et al. Genome-Wide Analysis of Cold Adaptation in Indigenous Siberian Populations. PLOS ONE 9, e98076 (2014).

20. Chrobok, L. et al. Timekeeping in the hindbrain: a multi-oscillatory circadian centre in the mouse dorsal vagal complex. Commun. Biol. 3, 1–12 (2020).

21. Vollebregt, M. A. et al. The Role of Gene Encoding Variation of DRD4 in the Relationship between Inattention and Seasonal Daylight. 825083 Preprint at https://doi.org/10.1101/825083 (2019).

22. Hwang, C. K. et al. Circadian Rhythm of Contrast Sensitivity Is Regulated by a Dopamine– Neuronal PAS-Domain Protein 2–Adenylyl Cyclase 1 Signaling Pathway in Retinal Ganglion Cells. J. Neurosci. 33, 14989–14997 (2013).

23. Jackson, C. R., Chaurasia, S. S., Hwang, C. K. & Iuvone, P. M. Dopamine D4 receptor activation controls circadian timing of the adenylyl cyclase 1/cyclic AMP signaling system in mouse retina. Eur. J. Neurosci. 34, 57–64 (2011).

24. Khazaal, A. Q. et al. Aryl hydrocarbon receptor affects circadian-regulated lipolysis through an E-Box-dependent mechanism. Mol. Cell. Endocrinol. 559, 111809 (2023).

25. Jaeger, C. & Tischkau, S. A. Role of Aryl Hydrocarbon Receptor in Circadian Clock Disruption and Metabolic Dysfunction. Environ. Health Insights 10, 133–141 (2016).

26. Griffin, P. et al. Circadian clock protein Rev-erbα regulates neuroinflammation. Proc. Natl. Acad. Sci. 116, 5102–5107 (2019).

27. Yokoyama, Y., Lambeck, K., De Deckker, P., Johnston, P. & Fifield, L. K. Timing of the Last Glacial Maximum from observed sea-level minima. Nature 406, 713–716 (2000).

28. Rankama, T. & Ukkonen, P. On the early history of the wild reindeer (Rangifer tarandus L.) in Finland. Boreas 30, 131–147 (2001).

29. Røed, K. H. et al. Historical and social–cultural processes as drivers for genetic structure in Nordic domestic reindeer. Ecol. Evol. 11, 8910–8922 (2021).

30. Røed, K. H., Bjørklund, I. & Olsen, B. J. From wild to domestic reindeer – Genetic evidence of a non-native origin of reindeer pastoralism in northern Fennoscandia. J. Archaeol. Sci. Rep. 19, 279–286 (2018).

31. Harding, L. E. Available names for Rangifer (Mammalia, Artiodactyla, Cervidae) species and subspecies. ZooKeys 1119, 117–151 (2022).

32. Røed, K. H., Kvie, K. S. & Bårdsen, B.-J. Genetic structure and origin of semi-domesticated reindeer. in Reindeer Husbandry and Global Environmental Change (Routledge, 2022).

33. Svishcheva, G. et al. Genetic differentiation between coexisting wild and domestic Reindeer (Rangifer tarandus L. 1758) in Northern Eurasia. Genet. Resour. 3, 1–14 (2022).

34. Kharzinova, V. et al. Insight into the Current Genetic Diversity and Population Structure of Domestic Reindeer (Rangifer tarandus) in Russia. Animals 10, 1309 (2020).

35. Lv, F.-H. et al. Whole-Genome Resequencing of Worldwide Wild and Domestic Sheep Elucidates Genetic Diversity, Introgression, and Agronomically Important Loci. Mol. Biol. Evol. 39, msab353 (2022).

36. Mei, C. et al. Genetic Architecture and Selection of Chinese Cattle Revealed by Whole Genome Resequencing. Mol. Biol. Evol. 35, 688–699 (2018).

37. Abri, M. A. A., Holl, H. M., Kalla, S. E., Sutter, N. B. & Brooks, S. A. Whole genome detection of sequence and structural polymorphism in six diverse horses. PLOS ONE 15, e0230899 (2020).

38. MacGillivray, D. M. & Kollmann, T. R. The Role of Environmental Factors in Modulating Immune Responses in Early Life. Front. Immunol. 5, 434 (2014).

39. Chouchani, E. T. & Kajimura, S. Metabolic adaptation and maladaptation in adipose tissue. Nat. Metab. 1, 189–200 (2019).

40. Jangam, D., Feschotte, C. & Betrán, E. Transposable element domestication as an adaptation to evolutionary conflicts. Trends Genet. TIG 33, 817–831 (2017).

41. Chessa, B. et al. Revealing the history of sheep domestication using retrovirus integrations. Science 324, 532–6 (2009).

42. Lieberman-Aiden, E. et al. Comprehensive Mapping of Long-Range Interactions Reveals Folding Principles of the Human Genome. Science 326, 289–293 (2009).

43. Zaharia, M., et al. Faster and More Accurate Sequence Alignment with SNAP. ArXiv11115572 Cs Q-Bio (2011).

44. Durand, N. C. et al. Juicer provides a one-click system for analyzing loop-resolution Hi-C experiments. Cell Syst. 3, 95–98 (2016).

45. Oliver, J. L., Carpena, P., Hackenberg, M. & Bernaola-Galván, P. IsoFinder: computational prediction of isochores in genome sequences. Nucleic Acids Res. 32, W287–292 (2004).

46. Smith, A. D., Sumazin, P., Xuan, Z. & Zhang, M. Q. DNA motifs in human and mouse proximal promoters predict tissue-specific expression. Proc. Natl. Acad. Sci. U. S. A. 103, 6275–6280 (2006).

47. Kerpedjiev, P. et al. HiGlass: web-based visual exploration and analysis of genome interaction maps. Genome Biol. 19, 125 (2018).

48. Tarailo-Graovac, M. & Chen, N. Using RepeatMasker to identify repetitive elements in genomic sequences. Curr. Protoc. Bioinforma. Chapter 4, Unit 4.10 (2009).

49. Stanke, M. et al. AUGUSTUS: ab initio prediction of alternative transcripts. Nucleic Acids Res. 34, W435–W439 (2006).

50. Korf, I. Gene finding in novel genomes. BMC Bioinformatics 5, 59 (2004).

51. Dobin, A. et al. STAR: ultrafast universal RNA-seq aligner. Bioinformatics 29, 15–21 (2013).

52. Cantarel, B. L. et al. MAKER: An easy-to-use annotation pipeline designed for emerging model organism genomes. Genome Res. 18, 188–196 (2008).

53. Holt, C. & Yandell, M. MAKER2: an annotation pipeline and genome-database management tool for second-generation genome projects. BMC Bioinformatics 12, 491 (2011).

54. Lowe, T. M. & Eddy, S. R. tRNAscan-SE: A Program for Improved Detection of Transfer RNA Genes in Genomic Sequence. Nucleic Acids Res. 25, 955–964 (1997).

55. Andrews, S. A quality control analysis tool for high throughput sequencing data: s-andrews/FastQC. (2019).

56. Ewels, P., Magnusson, M., Lundin, S. & Käller, M. MultiQC: summarize analysis results for multiple tools and samples in a single report. Bioinformatics 32, 3047–3048 (2016).

57. Li, H. & Durbin, R. Fast and accurate short read alignment with Burrows–Wheeler transform. Bioinformatics 25, 1754–1760 (2009).

58. Li, H. et al. The Sequence Alignment/Map format and SAMtools. Bioinforma. Oxf. Engl. 25, 2078–9 (2009).

59. Van der Auwera, G. A. et al. From FastQ data to high confidence variant calls: the Genome Analysis Toolkit best practices pipeline. Curr. Protoc. Bioinforma. 43, 11.10.1–11.10.33 (2013).

60. Narasimhan, V. et al. BCFtools/RoH: a hidden Markov model approach for detecting autozygosity from next-generation sequencing data. Bioinforma. Oxf. Engl. 32, 1749–1751 (2016).

61. R: A language and environment for statistical computing. (2019).

62. Zheng, X. et al. A high-performance computing toolset for relatedness and principal component analysis of SNP data. Bioinforma. Oxf. Engl. 28, 3326–3328 (2012).

63. Korneliussen, T. S., Albrechtsen, A. & Nielsen, R. ANGSD: Analysis of Next Generation Sequencing Data. BMC Bioinformatics 15, 356 (2014).

64. Fumagalli, M., Vieira, F. G., Linderoth, T. & Nielsen, R. ngsTools: methods for population genetics analyses from next-generation sequencing data. Bioinforma. Oxf. Engl. 30, 1486–1487 (2014).

65. Pfeifer, B., Wittelsbürger, U., Ramos-Onsins, S. E. & Lercher, M. J. PopGenome: An Efficient Swiss Army Knife for Population Genomic Analyses in R. Mol. Biol. Evol. 31, 1929–1936 (2014).

66. Stajich, J. E. et al. The Bioperl toolkit: Perl modules for the life sciences. Genome Res. 12, 1611– 1618 (2002).

67. Lee, T.-H., Guo, H., Wang, X., Kim, C. & Paterson, A. H. SNPhylo: a pipeline to construct a phylogenetic tree from huge SNP data. BMC Genomics 15, 162 (2014).

68. Felsenstein, J. PHYLIP (Phylogeny Inference Package) version 3.6. Distributed by the author. Department of Genome Sciences, University of Washington, Seattle. (2005).

69. Rambaut, A. FigTree. (2018).

70. Danecek, P. et al. The variant call format and VCFtools. Bioinformatics 27, 2156–2158 (2011).

71. Kumar, S., Stecher, G. & Tamura, K. MEGA7: Molecular Evolutionary Genetics Analysis Version 7.0 for Bigger Datasets. Mol. Biol. Evol. 33, 1870–1874 (2016).

72. Alexander, D. H., Novembre, J. & Lange, K. Fast model-based estimation of ancestry in unrelated individuals. Genome Res. 19, 1655–1664 (2009).

73. Purcell, S. et al. PLINK: a tool set for whole-genome association and population-based linkage analyses. Am. J. Hum. Genet. 81, 559–75 (2007).

74. Francis, R. M. pophelper: an R package and web app to analyse and visualize population structure. Mol. Ecol. Resour. 17, 27–32 (2017).

75. Quinlan, A. R. & Hall, I. M. BEDTools: A Flexible Suite of Utilities for Comparing Genomic Features. Bioinformatics 26, 841–2 (2010).

76. Mailund, T. et al. A New Isolation with Migration Model along Complete Genomes Infers Very Different Divergence Processes among Closely Related Great Ape Species. PLoS Genet. 8, e1003125 (2012).

77. Shen, W., Le, S., Li, Y. & Hu, F. SeqKit: A Cross-Platform and Ultrafast Toolkit for FASTA/Q File Manipulation. PLOS ONE 11, e0163962 (2016).

78. Kumar, S. & Subramanian, S. Mutation rates in mammalian genomes. Proc. Natl. Acad. Sci. U. S. A. 99, 803–808 (2002).

