## Supplementary information for "Whole genome sequencing provides novel insights into the evolutionary history and genetic adaptations of reindeer populations in northern Eurasia"

Supplementary figure legends

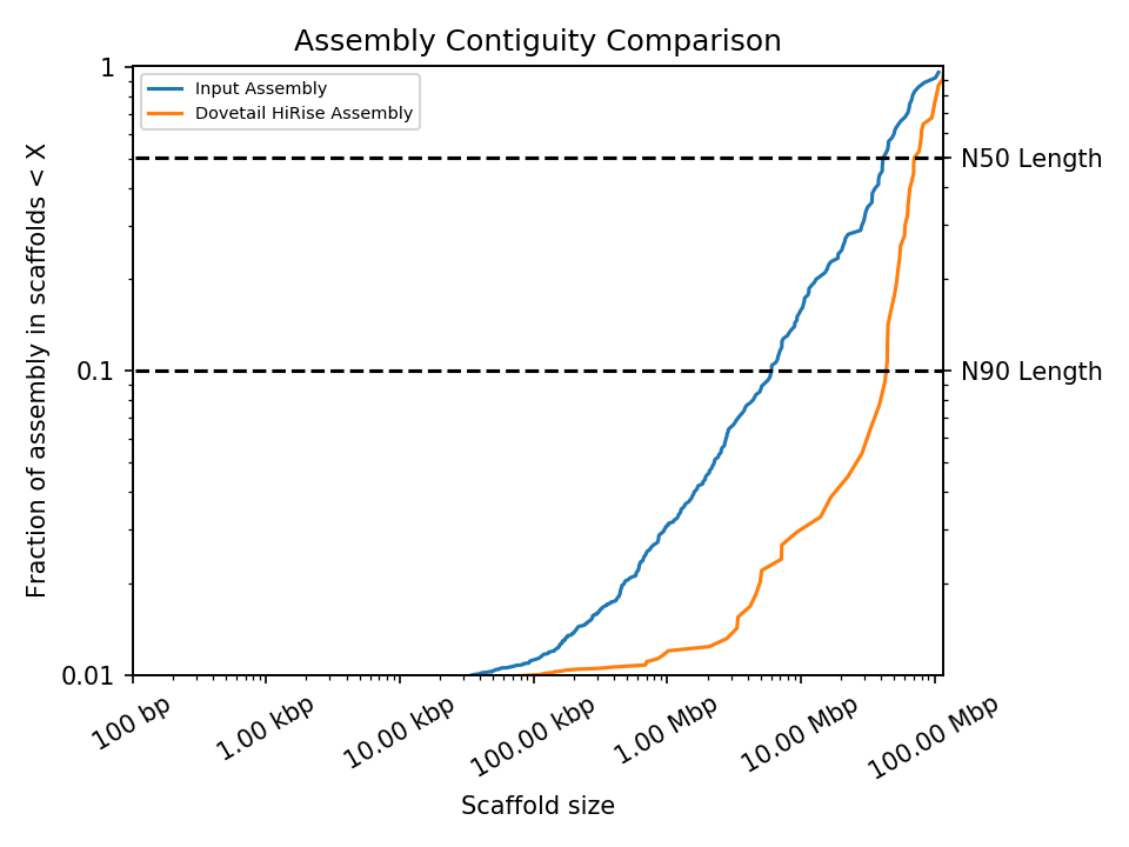

Supplementary Figure 1: Contiguity of current reindeer assembly. A comparison of the contiguity of the input assembly and the final HiRise scaffolds. Each curve shows the fraction of the total length of the assembly present in scaffolds of a given length or smaller. The fraction of the assembly is indicated on the Y-axis and the scaffold length in basepairs is given on the X-axis. The two dashed lines mark the N50 and N90 lengths of each assembly. Scaffolds less than 1 kb are excluded.

 
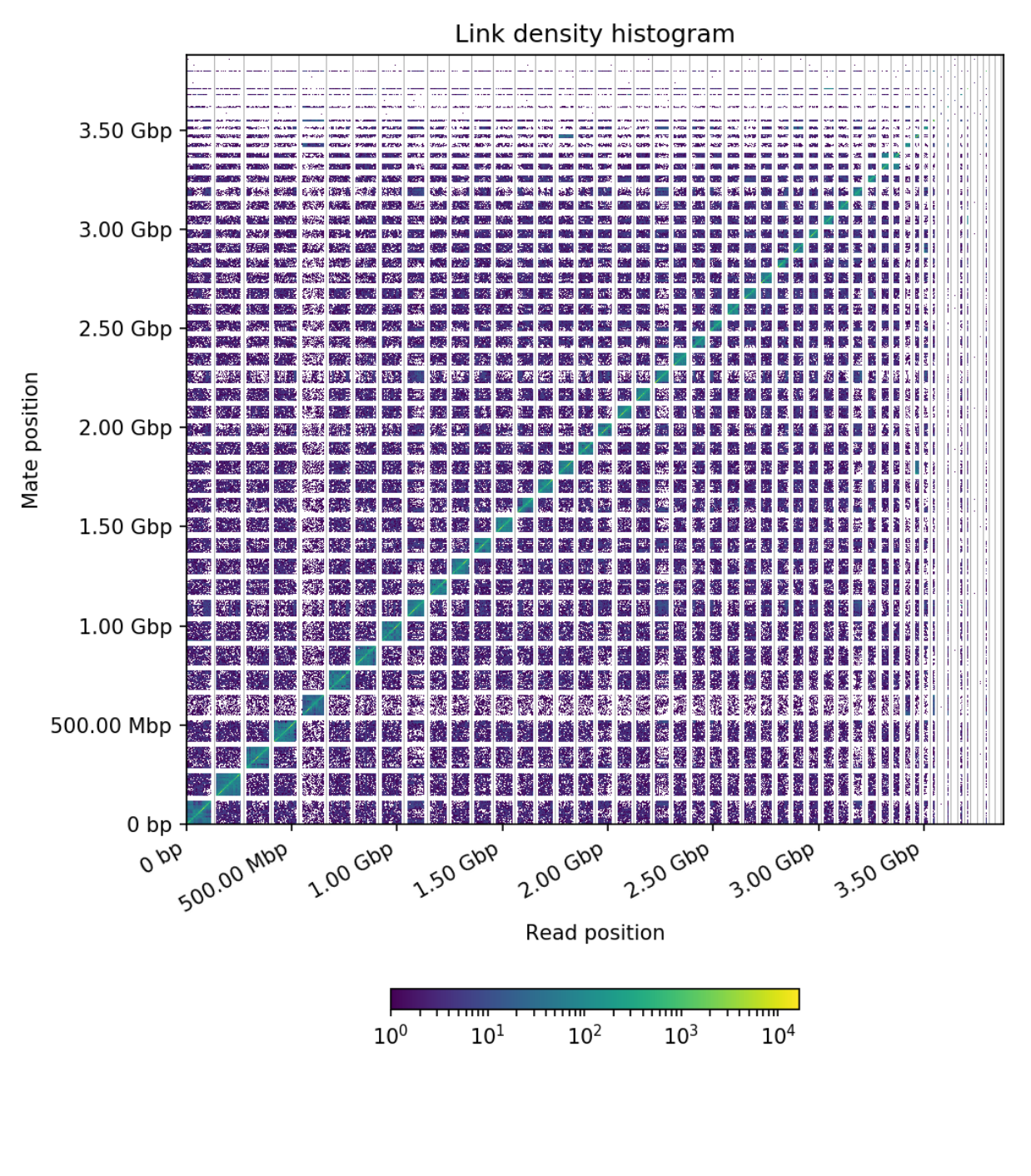

Supplementary Figure 2: Link density histogram of reindeer assembly. In this figure, the x and y axes give the mapping positions of the first and second read in the read pair respectively, grouped into bins. The color of each square gives the number of read pairs within that bin. White vertical and black horizontal lines have been added to show the borders between scaffolds. Scaffolds less than 1 Mb are excluded.

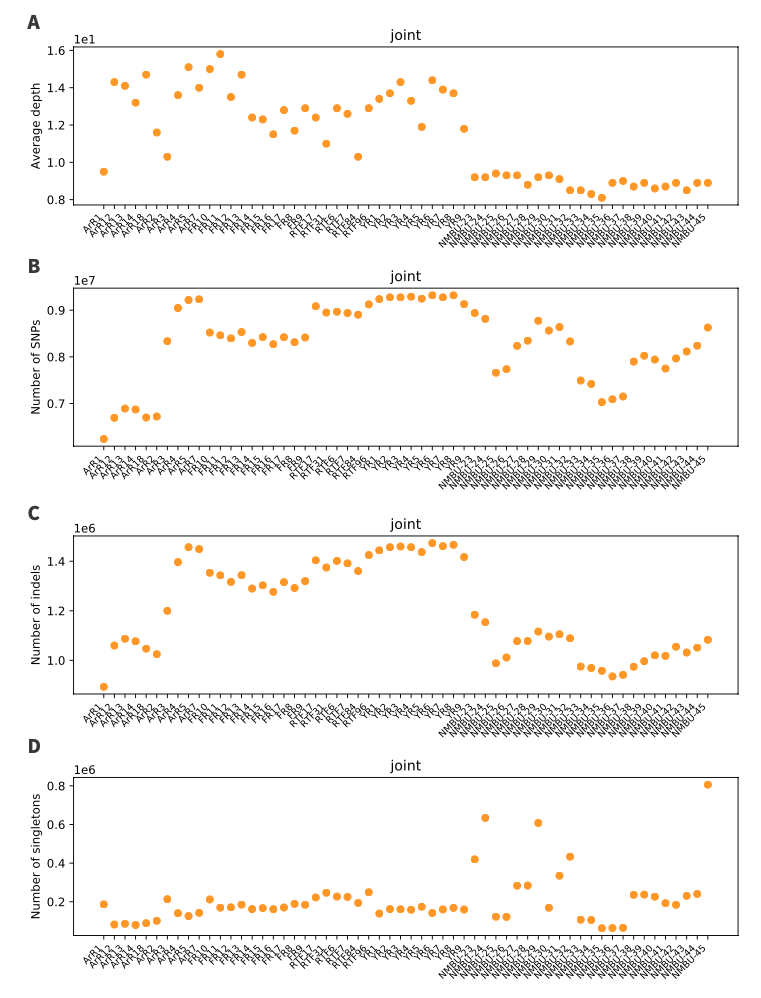

Supplementary Figure 3: Plots of sample-wise variant call statistics, showing average sequencing depth at variant sites (A), number of SNPs (B), number of indels (C) and number of singletons (D) per sample.

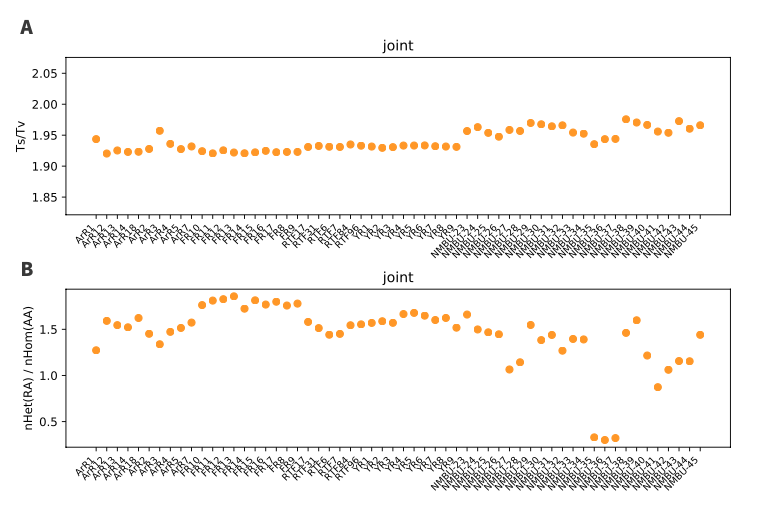

Supplementary Figure 4: Plots of sample-wise transition vs transversion (Ts/Tv) ratios (A) and heterozygous vs homozygous SNP ratios (B).

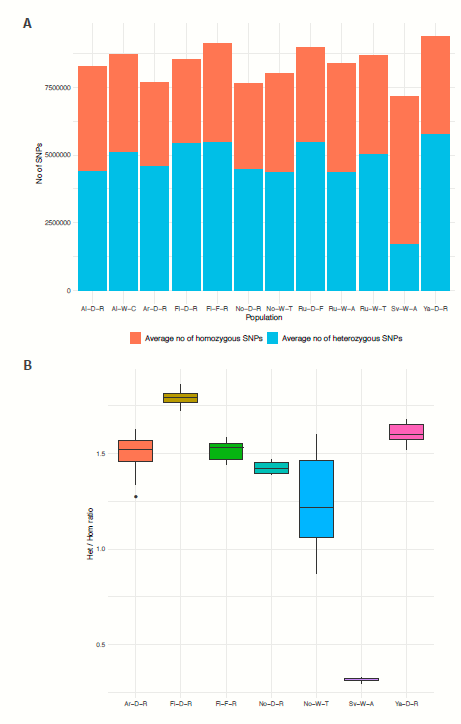
remove Figure S5A

Supplementary Figure 5: (A)Average SNP count per individual within each population, with the proportions of homozygous and heterozygous SNPs are shown in red and blue, respectively.(B) Distributions of the ratios of heterozygous to homozygous SNPs within seven main populations.

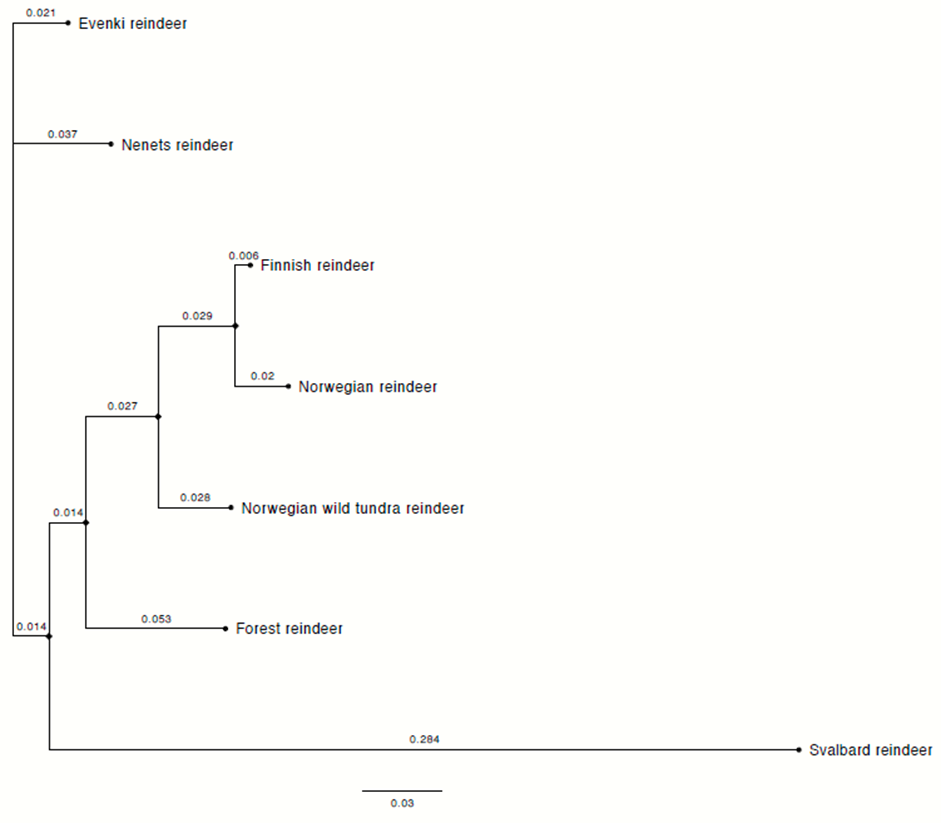

Supplementary Figure 6. Neighbour-joining tree constructed based on the pairwise FST values of the seven main populations selected based on the PCA. The population details are described in Table 2. The branch lengths are indicated on the branches. pairwise FST values of the seven main populations selected based on the PCA. The population details are described in Table 2. The branch lengths are indicated on the branches.

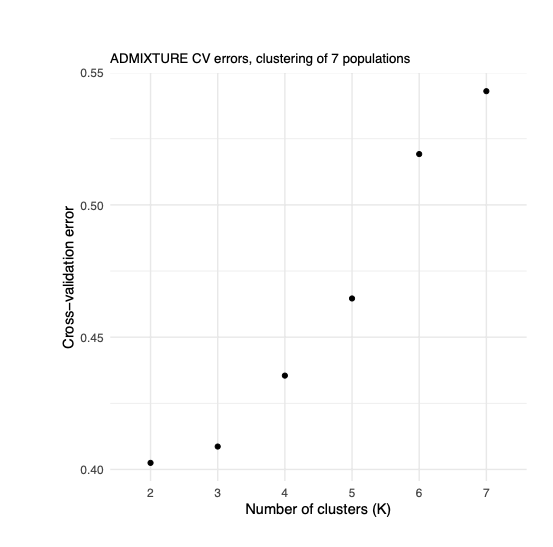

Supplementary Figure 7. ADMIXTURE 5-fold cross-validation errors for the population structure analysis of the seven main populations. K values ranging from 2 to 7 were tested.

Supplementary tables:

Supplementary Table 1: TAD statistics

| Resolution (kbp) | Number of TADs | Mean TAD Size (bp) | Basepairs in TADs (kbp) | Percent genome contained in TADs |
| --- | --- | --- | --- | --- |
| 10 | 263 | 645,741 | 1,666 | 6.44% |
| 25 | 1,491 | 655,549 | 9,266 | 35.84% |
| 50 | 1,452 | 1,339,325 | 15,307 | 59.2% |

Supplementary Table 2: Summary table of the main population diversity statistics (pi and FST) of the seven main populations

| Population | Nucleotide diversity (pi) within population | Watterson’s theta |
| --- | --- | --- |
| Fi-D-R | 0.000642 | 0.000650 |
| No-D-R | 0.000602 | 0.000592 |
| No-W-T | 0.000642 | 0.000635 |
| Fi-F-R | 0.000650 | 0.000633 |
| Ar-D-R | 0.000650 | 0.000622 |
| Ya-D-R | 0.000669 | 0.000661 |
| Sv-W-A | 0.000231 | 0.000217 |

Supplementary Table 3: Pairwise F_ST_ values of the seven main populations: Finnish semi-domestic reindeer (Fi-D-R), Norwegian semi-domestic reindeer (No-D-R), Norwegian wild tundra reindeer (No-W-T), Finnish wild forest reindeer (Fi-F-R), Nenets semi-domestic reindeer (Ar-D-R) and Even semi-domestic reindeer (Ya-D-R).

|  | Ar-D-R | Fi-D-R | Fi-F-R | No-D-R | No-W-T | Sv-W-A |
| --- | --- | --- | --- | --- | --- | --- |
| Fi-D-R | 0.110104 |  |  |  |  |  |
| Fi-F-R | 0.113016 | 0.098953 |  |  |  |  |
| No-D-R | 0.128794 | 0.009113 | 0.121493 |  |  |  |
| No-W-T | 0.114427 | 0.064951 | 0.102973 | 0.071922 |  |  |
| Sc-W-A | 0.402246 | 0.403715 | 0.399765 | 0.424133 | 0.397652 |  |
| Ya-D-R | 0.056069 | 0.099039 | 0.101648 | 0.117213 | 0.101618 | 0.388949 |
